# Molecular identification and functional analysis of HrpZ2, a new member of harpin superfamily from *Pseudomonas syringae* inducing hypersensitive response in tobacco

**DOI:** 10.1101/2025.07.14.664668

**Authors:** Kishori Lal, Anurag Joshi, Vartika Saini, Mujahid Mohammed, P.V.S.R.N Sarma, Debashish Dey

## Abstract

Harpins belong to a group of proteins with distinctive features such as heat stability, glycine richness, and absence of cysteine, and are secreted by many Gram-negative phytopathogens *via* the type III secretion system. Harpins are known to trigger hypersensitive response followed by induction of systemic acquired resistance in non-host plants. However, the molecular mechanism of harpin-induced hypersensitive response remained largely unexplored, mainly because of lack of structural information. In this study, we report the cloning of a new harpin gene (*hrpZ2*) from *Pseudomonas syringae* strain MTCC-11950, belonging to the harpin superfamily. *In silico* analysis revealed that about 50.29% of the protein consists of alpha-helices, 48.53% are random coils, and only 1.16% are beta-sheets, and nearly half (42%) of the protein consists of intrinsically disordered regions. Based on a prokaryotic predictive model and the presence of a signal peptide on its N-terminus, the subcellular localization of harpin is predicted as extracellular. To date, no experimentally determined crystal structure of any harpin protein is available. Therefore, we built and validated a three-dimentional model (with 99% of residues in allowed/additionally allowed regions and a Z-score of −5.3) of harpin. Phylogenetic analysis and functional domain studies revealed that this new harpin belongs to the harpin superfamily. Infiltration of harpin in tobacco leaves resulted in hypersensitive response, which was associated with oxidative burst, callose deposition, localized cell death, and increased activity of defense-related enzymes such as phenylalanine ammonia-lyase and polyphenol oxidase. Furthermore, infiltration of harpin in non-host plants from different angiosperm families induced hypersensitive response, indicating broad-spectrum agricultural applicability of this new harpin protein. This study elucidates the molecular and functional properties of the new harpin protein and its ability to induce hypersensitive response across a broad range of non-host plants.

## 1 Introduction

The interactions between plants and plant pathogens are shaped by a dynamic and continuous co-evolutionary process. Plant pathogens deploy diverse strategies to suppress host defenses and establish infection, while plants, in turn, have evolved a multilayered immune system to detect and restrict invading pathogens (Jones and Dangl, 2006; Cui et al., 2015). This innate immune system is broadly categorized into two hierarchical layers: PAMP triggered immunity (PTI) and effector triggered immunity (ETI).

The first layer, PTI, involves the recognition of the pathogen’s surface molecular patterns known as pathogen-associated molecular patterns (PAMP) by plant plasma membrane-bound receptors defined as pattern recognition receptors (PRR). Usually, PAMPs are considered to be essential for general microbial fitness and survival. These PAMP-PRR complexes activate the immune responses termed as PAMP-triggered immunity (PTI) in plants (Zipfel et al., 2006; Dodds and Rathjen, 2010). However, adapted pathogens have evolved the capacity to overcome PTI-mediated immunity through deployment of effector proteins into the host cytoplasm *via* specialized secretion systems, notably the Type-III secretion system (T3SS) in Gram negative bacteria resulting in effector-triggered susceptibility (ETS) (Martel et al., 2021). These effectors target the host signaling and contribute to the virulence of invading pathogens. On the other side, plants got evolved parallelly and acquired more receptors that specifically recognize and bind to the pathogen effectors and mount a second layer of defense response known as effector-triggered immunity (ETI). The ETI leads to an immediate oxidative burst (ROS) followed by the hypersensitive response (HR) mediated cell death of plant cells at and near the sites of pathogen invasion. HR was first time described by Wei et al. (1992), as a rapid, highly localized defense response that is generally considered as a form of programmed cell death (PCD) in plants. HR is often followed by systemic acquired resistance (SAR), a salicylic acid (SA)-dependent defense response throughout the plant which is effective against a broad-spectrum of phytopathogens (Ricci, 1997).

Elicitors derived from microbial sources may function as molecular signals that trigger a myriad of innate immune responses such as oxidative burst, HR-mediated programmed cell death, accumulation of callose, lignification, pathogenesis-related (PR) proteins, and the activation of key defense enzymes like polyphenol oxidase (PPO) and phenylalanine ammonia-lyase (PAL). Thus, these elicitors offer a promising environmentally sustainable alternative to traditional chemical pesticides (Choi et al., 2013; Zhu et al., 2024).

Certain Gram-negative, plant-pathogenic bacteria harbor pathogenicity (*hrp*) gene clusters that regulate HR in nonhost plants or resistant cultivars of host plants and control the pathogenicity in susceptible plants (Collmer et al., 1998). Generally, the *hrp* cluster contains genes that encode the components of a bacterial type III secretion system, through which the effector proteins are injected and delivered into the host’s cytosol (Galán and Collmer, 1999). Harpin (*hrpN*) was the first bacterial elicitor protein identified by Wei et al., (1992) from the *Erwinia amylovora* and named as Harpin*_Ea_*. To date, several other *hrp*-encoded harpin elicitors have been identified from various phytopathogenic bacteria collectively termed as the Harpin protein family (Choi et al., 2013). Recently, all these harpins were classified into five major groups based on their inherent domain structure and protein similarity, *viz*., HrpN group (from *Erwinia* spp.), HrpZ1 group (from *P. syringae*), HrpW1 (from *Erwinia* spp., *P. syringae*, *Ralstonia* spp.), Hpa1 group and its orthologs (from *Xanthomonas* spp.), and ‘Others’ group contained remaining unclassified harpin proteins (Choi et al., 2013). The HrpZ-group of harpins are a major T3SS-dependent proteins secreted by *P. syringae* which shares some common characteristics with other groups of harpins: such as being glycine-rich, lack of cysteine (exception Hpa1), very few aromatic amino acids and high thermal stability (Choi et al., 2013; Tarafdar et al., 2009). A recent study on the thermal stability of harpins showed that the harpin (HpaXpm) remains active at 100°C, 150°C, or 200°C while the ability of heat-treated harpins to HR induction varied with respect to temperature (Liu et al., 2020). The reason behind the high thermal stability of harpin proteins might be due to their inherent structural features rendering thermostability. It is still unknown and yet to be explored (Tarafdar et al., 2009). Harpins can act as an elicitor to elicit an HR associated with systemic acquired resistance (SAR) and provide diverse benefits to plants, such as broad-spectrum durable disease resistance, quality, and yield improvement in crops. Harpin (HrpN) protein from *E. amylovora*, enhanced drought resistance in *Arabidopsis thaliana* by activating the abscisic acid (ABA)-dependent signalling pathways (Dong et al., 2005). Similarly, harpin (HrpZ) from *Pseudomonas* spp., has been reported to increase disease resistance in sugar beet (*Beta vulgaris*) and *Nicotiana benthamiana* against rhizomania disease caused by Beet necrotic yellow vein virus (Pavli et al., 2011). Moreover, harpin proteins have also been involved in ethylene-mediated resistance against insect herbivory and promoting overall plant growth and vigour (Dong et al., 2004). Recent reports (Li et al., 2015) showed that aquaporin proteins present on the plasma membranes act as receptors for the perception of extracellular harpin*_Xoo_* proteins. A recent report from our laboratory identified a signature sequence present in the aquaporin protein and the candidate amino acid residues involved in the harpin-aquaporin interaction through *in silico* docking studies (Lal et al., 2023).

These findings demonstrate that harpins elicit diverse physiological and defense responses in non-host plants. Given their potential to enhance tolerance to both biotic and abiotic stresses, harpins are a promising tool for sustainable crop management. To realize their full potential, it is essential to identify and screen new harpin variants with superior elicitor activity and broader applicability for crop improvement.

In this study, we present the molecular and functional characterization of HrpZ2*_Ps_*, a newly identified member of the harpin superfamily of effectors from *P. syringae*. The harpin gene encodes a 34.5 kDa predicted protein, exhibiting characteristic features of harpins, including a glycine-rich region, and a potential secondary structure propensity. Functional assays evaluated its capacity to elicit HR in diverse non-host plants, and activate defense-related enzymes in tobacco, thereby asserting its broad-spectrum disease resistance potential for agricultural applications.

## 2 Materials and methods

### 2.1 Plants, bacterial strains, and growth conditions

The seeds of tobacco plant (*Nicotiana tabacum* cv. Xanthi) were maintained at 4°C in the Laboratory of Plant Biotechnology. For this study, seeds were sown in flat plastic trays (32 x 15 cm) filled with a 1:1:1(v/v/v) mixture of vermiculite, garden soil, and sand. At the four-leafed stage, seedlings were transplanted individually into 12 x 10 cm plastic pots (one plant per pot). Plants were maintained at 23 ± 3°C, under a 16 h light and 8 h dark photoperiod.

We procured five different strains of *P. syringae* from the bacterial culture collections of MTCC, Chandigarh, India, with the aim of cloning new harpin genes. The *P. syringae* strains were cultured on Luria-Bertani (LB) Agar (Himedia, M1151) and Luria-Bertani (LB) broth (Himedia, M1245) at 28°C. *Escherichia coli* DH5α cells (used for gene cloning) and BL21 cells (used for expression studies) were cultured on LB Agar and LB broth at 37°C. Kanamycin (50 µg/mL) was used for bacterial selection.

### 2.2 NCBI Submission of the harpin gene

We used previously reported harpin oligo pairs (forward and reverse) (Madhuri et al., 2012; Dey et al., 2014) in PCR reaction for new harpin gene amplification from five different strains of *P. syringae*. We could successfully amplify a band of 1.02 kb, corresponding to the harpin gene from only one strain (MTCC-11950) out of the five bacterial strains, which was cloned and sequenced. DNA sequence analysis revealed that it is a previously uncharacterized member of the harpin protein superfamily, designated as *hrpZ2*. The sequence has been deposited in the National Center for Biotechnology information (NCBI) GenBank (accession ID: OQ338148.1).

### 2.3 *In silico* analysis of the primary, secondary and tertiary structures of harpin

#### 2.3.1 Sequence retrieval and sequence-based analysis

The amino acid sequence of harpin (HrpZ2*_Ps_*) was retrieved from the NCBI GenBank: OQ338148.1, in the FASTA format. The physiochemical properties of harpin were analyzed using the ProtParam tool (https://web.expasy.org/protparam/). Hydropathicity or hydrophobicity, isoelectric point (pI), instability index and molecular weight of harpin sequence were analyzed using the Grand Average of Hydropathicity (GRAVY) web server. Intrinsic disorder was evaluated with the PrDOS server (https://prdos.hgc.jp/cgi-bin/top.cgi) (Ishida and Kinoshita, 2007).

#### 2.3.2 Three-dimentional (3D) modelling of harpin

Determination of protein sequence and structure is fundamental to understanding its biological function. Currently, there is no experimentally concluded tertiary structure available for any harpin in the Protein Data Bank (PDB). Consequently, the tertiary structure of the HrpZ2*_Ps_* protein (GenBank ID: WDY61303.1) was modeled by utilizing the I-TASSER server (https://zhanggroup.org/I-TASSER/) (Zheng et al., 2025). Tools such as VERIFY 3D (Colovos and Yeates, 1993), ERRAT (Bowie, J U et al., 1991), and PROCHECK (Laskowski et al., 1996) of the SAVES v.6.0 server (https://saves.mbi.ucla.edu) were used to analyze the Ramachandran Plot and validate the computationally predicted 3D structure. Z-score was calculated using the ProSA-web server (https://prosa.services.came.sbg.ac.at/prosa.php).

#### 2.3.3 Functional domain prediction and analysis of domain architecture

Conserve domains (CD) analysis was performed using NCBI’s CD search tool (https://www.ncbi.nlm.nih.gov). Harpin domains were further detected and annotated with the PSIPRED server (https://bioinf.cs.ucl.ac.uk/psipred) (Cozzetto et al., 2016), which infers functional domains from the predicted secondary structure. The transmembrane helix packing and orientations were evaluated *via* the PSIPRED workbench using the MEMPACK alpha-helical transmembrane protein structure prediction server (Nugent et al., 2011). Helix and loop regions were predicted with MEMSAT-SVM algorithms, and their orientations were cross-validated with the TMHMM web server and the kyte-doolittle hydropathy profiling method of the PSIPRED Workbench (Nugent and Jones, 2009).

#### 2.3.4 Evolutionary study of Harpin

The evolutionary history was inferred using the Maximum Likelihood (ML) method, and the phylogenetic tree was constructed using MEGA11.0 software (https://www.megasoftware.net/older_versions). Bootstrap analysis with 1000 replicates was performed to ensure statistical robustness and reliability of the branching patterns. The phylogenetic analysis was done among 23 harpin proteins derived from different bacterial genera, such as *Xanthomonas, Pseudomonas, Ralstonia,* and *Erwinia.* The phylogenetic tree was visualized on the iTOL online tool (https://itol.embl.de/shared/2PuqbCYlUfiSH) (Letunic and Bork, 2024).

#### 2.3.5 Subcellular localization of harpin

The subcellular localization of HrpZ2*_Ps_* was predicted using DeepLocPro - 1.0 (https://services.healthtech.dtu.dk/services/DeepLocPro-1.0/). The server predicted the subcellular localizations using the prokaryotic predictive model. It provides the prediction probability for different subcellular compartments like outer membrane, periplasmic space, cell wall and surface, cytoplasmic membranes based on the amino acid sequence of the signal sequence of the protein.

### 2.4 Cloning, expression, and purification of harpin

To construct recombinant plasmids encoding poly-histidine-tailed harpin fusion protein, initially, the harpin gene was amplified from the genomic DNA of *P. syringae* strain (MTCC-11950) using a pair of primers: 5’-CGAATCCCATATGCAGAGTCTCAGTCTTAAC – 3’ (forward primer) and 5’-AGGATCCTCGAGGGCTGCAGCCTGATTGC – 3’ (reverse primer). The harpin gene (*hrpZ2*, 1.02 kb) encoding harpin protein was cloned in the *pET28a* vector (maintained in our laboratory) under the *Nde*I and *Xho*I restriction sites. The recombinant plasmids (*pET28a*-*hrpZ2*) were transformed into an *E. coli* BL21 (DE3) expression host and grown in LB broth supplemented with Kanamycin (50 µg/ml) till the OD_600nm_∼ 0.5 and the expression of harpin protein was induced with 1 mM Isopropyl-β-D-thiogalactopyranoside (IPTG) at 37°C for 3 h with shaking conditions (200 rpm). The bacterial cells were harvested using centrifugation at 4,830 rcf at 4°C and the pellet was resuspended in 10 mM sodium phosphate buffer (pH 7.5), supplemented with 1 mg/ml lysozyme and immediately sonicated using probe sonicator (Labman, 3 mm probe) using pulse rate of 30 s and 20 s on/off respectively, with 40% amplitude, for 20 minutes with two minutes interval to cool down. The sonicated lysate was centrifuged at 13,528 rcf for 30 min and the supernatant was loaded onto a nickel-nitrilotriacetic acid (Ni-NTA, Qiagen, USA) affinity chromatography column filled with Ni-NTA slurry. The protein was purified following the manufacturer’s protocol (Manual purification of 6xHis-tagged proteins from *E. coli*, Qiagen) with some modifications. The targeted, 6xHis-tagged harpin proteins got bound to the nickel in the affinity column. Subsequently, the column was washed with a washing buffer containing 10 mM sodium phosphate buffer (pH 7.5) supplemented with 20 mM and 30 mM imidazole to remove nonspecifically bound or loosely attached proteins. Finally, the targeted harpin protein was eluted out using an elution buffer consisting of 10 mM sodium phosphate buffer (pH 7.5) with 200 mM imidazole. The purified protein was extensively dialyzed using dialysis membrane tubing (10 kDa cut off, SnakeSkin™ Dialysis Tubing, Thermo Scientific™, USA) against 10 mM sodium phosphate buffer (pH 7.5) without imidazole. The dialyzed protein was concentrated using an Amicon centrifuge column (10 kDa cut-off, Millipore, USA). The purity of the protein was checked by running on a 12% SDS-PAGE gel. The estimation of protein concentration was done by following standard Bradford’s method (Bradford, 1976).

### 2.5 SDS-PAGE and western blotting of harpin (HrpZ2*_Ps_*)

For western blotting, the protein (HrpZ2*_Ps_*) was resolved on a 12% SDS-PAGE gel under reducing conditions along with a molecular weight marker and then transferred to a polyvinylidene difluoride (PVDF) membrane. The membrane was blocked with a blocking buffer containing 5% (w/v) non-fat milk in Tris Buffered Saline with Tween 20 (TBS, 150 mM NaCl and 0.05% Tween 20, pH 7.4) at room temperature for one hour, followed by incubation with harpin primary anti-His-tag antibody in rabbit (GenScript, A00174), diluted 1:5000 in 1x TBST with 5% milk, incubated at 4°C with gentle shaking for overnight. Further, the PVDF membrane was washed five times with TBST for 10 min each time. The membrane was then incubated with goat anti-rabbit IgG-ALP conjugated secondary antibody (Bangalore GeNei™, 1100180011730) in TBST (1:2000) at room temperature for 1 h and washed five times with TBST for 10 min each time. Finally, the immunoblot was visualized on the PVDF membrane using BCIP-NBT substrate for alkaline phosphatase.

### 2.6 Determination of the effective concentration of harpin protein to elicit HR in tobacco

Tobacco plants (*Nicotiana tabacum* cv. Xanthi) were grown under laboratory conditions at 23 ± 3°C, with a 16 h light and 8 h dark photoperiod (Dey et al., 2014). Five-week-old plants were used for infiltration with purified harpin (HrpZ2*_Ps_*). To assess the elicitation of HR, the HrpZ2*_Ps_* was diluted in 10 mM sodium phosphate buffer (pH 7.5) to a final concentration (30 µg/mL). Infiltration was performed on the abaxial surface of the third to fifth leaves from the apex using a 1 mL sterile needleless syringe. Buffer alone (10 mM Sodium phosphate buffer, pH 7.5) served as the negative control. Serial dilutions of purified harpin protein (1, 10, 20, 30, 40, 50 and 60 µg/ml) were also similarly infiltrated into the abaxial surface of leaves of tobacco to identify the effective concentration required for the induction of a visible macroscopic HR.

### 2.7 Host range determination of harpin protein using HR-induction assay

Harpins are proteinaceous elicitors well documented for their role in inducing plant defense responses, particularly through the elicitation of the hypersensitive response (HR). To evaluate the host-range of this new member of harpin elicitor derived from *P. syringae*, we assessed its ability to trigger localized HR in various angiosperm plant families. Affinity purified HrpZ2*_Ps_* was diluted in 10 mM sodium phosphate buffer (pH 7.5) to a final concentration of 50 µg/mL and infiltrated into the abaxial (lower) surface of the leaves of 12 non-host plants belonging to eight angiosperm plant families. A sterile needless syringe (1 mL) was used for the infiltration. Each leaf was infiltrated on one half with the purified harpin protein, while the opposite half was infiltrated with sodium phosphate buffer (10 mM, pH 7.5) as a negative control. Leaves were observed and photographed at different time intervals: before infiltration, and at 0, 12, 18, 24, 30 and 48 h post infiltration. A final observation and a photograph were taken five days post infiltration.

### 2.8 Biochemical assays

#### 2.8.1 Detection of reactive oxygen species (H_2_O_2_)

Hydrogen peroxide (H_2_O_2_) is a key signal molecule in the plant defense pathway, and its tissue accumulation can be visualized histochemically with 3, 3’-diaminobenzidine dyes (DAB). Purified harpin protein was syringe infiltrated into the interveinal region of the abaxial side of fully extended tobacco leaves using 1 mL sterile needleless syringe (final harpin concentration of 30 μg/ml) and the H_2_O_2_ detection was performed following Daudi and O’Brien, 2012 with some modifications. The third to fourth leaves were used for the infiltration. Sodium phosphate buffer (10 mM, pH 7.5) served as the negative control. For the detection of H_2_O_2_ accumulation, a 1 mg/ml DAB (Himedia, RM2735) solution was prepared in 10 mM sodium phosphate (Na_2_HPO_4_) buffer with 0.05% (v/v) Tween 20, adjusted to pH 3.8 with conc. HCL. The harpin treated leaves were harvested at 6-, 12-, and 24-hours post infiltration (hpi) and soaked in DAB-HCL solution (pH 3.8) for 10 hours at room temperature in dark conditions followed by boiling in bleaching solution (ethanol: acetic acid: glycerol in a 3:1:1, v/v/v ratio) for 15 minutes. Chlorophyll was fully bleached and the red-brown precipitate resulting from DAB’s reaction with H_2_O_2_ was visualized under light microscope and photographed. Each experiment was repeated three times, with six leaves from different plants per replicate. ROS signals were assessed in harpin treated tobacco leaf tissue by submerging in 10 µM 2′,7′–dichlorofluorescin diacetate (DCFDA, SIGMA-Aldrich, D6883-50MG) prepared in 10 mM sodium phosphate buffer (pH 7.5) and incubated in dark for 20 min at room temperature. Tissues were then gently washed with the same buffer, mounted on a glass slide and ROS signals were observed using a fluorescence microscope (Nikon, Japan).

#### 2.8.2 Quantification of Hydrogen peroxide using FOX method

Quantification of H_2_O_2_ was done using the FOX (Ferrous oxidation in xylenol orange) method as per Jiang et al., (1990) with few modifications. The absorbance (560 nm) of a coloured complex formed due to the oxidation of Fe^2+^ to Fe^3+^ by H_2_O_2_ and binding of the resulting Fe^3+^ to xylenol orange was measured. Briefly, the FOX reagent (250 μM ferrous ammonium sulfate (NH_4_)_2_Fe(SO_4_)_2_.6H_2_O, 100 μM xylenol orange, 25 mM H_2_SO_4_, and 100 mM sorbitol) was prepared by taking about 80 mL of 25 mM H_2_SO_4_, 9.8 mg ammonium ferrous sulfate, 7.6 mg xylenol orange, and 1.82 g sorbitol, mixed thoroughly until the sorbitol got completely dissolved, and the volume was increased to 100 mL with 25 mM H_2_SO_4_. At 0, 3, 6, 9, 12, 24, 48 and 72-hour post-infiltration (hpi), 0.5 g of leaf tissue were ground in liquid nitrogen and suspended in 5 mL ice-cold 0.1% trichloroacetic acid (TCA). The supernatant was separated by centrifugation at 18,894 rcf for 25 min at 4°C, 0.5 ml of supernatant was mixed with 0.5 ml of FOX reagent and incubated in dark for 30 minutes at room temperature. Absorbance was measured at 560 nm, using H_2_O_2_ as standard. Results were expressed in μM H_2_O_2_ per gram fresh weight (μM/g FW).

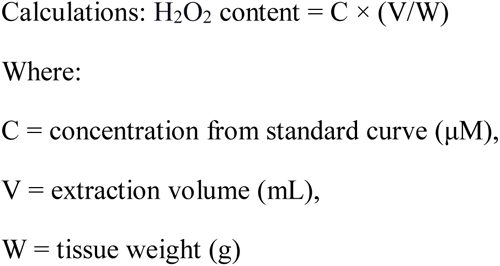

#### 2.8.3 Cell death estimation by using trypan blue staining

For the detection of dead cells in harpin treated tobacco leaves, trypan blue staining was performed as described by Koch and Slusarenko, 1990. The leaves were detached at 24, 48 and 72 hpi and boiled in lactophenol trypan blue solution containing (lactic acid (10 mL), glycerol (10 mL), trypan blue (7.6 mg) and phenol (10 ml) mixed in 10 mL distilled water). Full (uncut) leaves were boiled for approximately 1 min in the staining solution and then decolorized by using clearing solution (mixing 3 parts ethanol (96%) with 1-part acetic acid) (For 100 mL solution: 75 mL ethanol (96%), 25 mL glacial acetic acid). The leaves were stained at room temperature for 3-4 hours; the solution was replaced three times until the tissue became transparent. The de-stained leaves were rinsed with distilled water, followed by mounting in 60% glycerol and covered with a coverslip and carefully photographed using a light microscope (Magnus Opto Systems India Pvt. Ltd.).

#### 2.8.4 Detection of callose deposition

Callose accumulation was detected as per Luna et al., (2011) with some minor modifications. Tobacco plants at the six to eight-leaf-stage were syringe infiltrated onto the abaxial surface without a hypodermic needle. At 24-hpi, infiltrated leaf tissue was excised followed by chlorophyll removal, and the samples were completely dehydrated in 96% ethanol by three successive immersions. Specimens were briefly rinsed with distilled water, then equilibrated for 30 min in 10 mM sodium phosphate buffer (pH 7.5) at room temperature. Tissues were stained with 0.05% aniline blue for 1 h in dark at room temperature, mounted in 50% glycerol, and covered with a coverslip. Imaging was performed on a confocal laser scanning microscope (Zeiss, LSM780) using 405 nm excitation and 530 nm emission wavelength. Buffer alone (10 mM sodium phosphate, pH 7.5) served as a control. Callose intensity was quantified (using Image J software) by measuring the blue pixels on the image and the total pixels covering the plant material. Values were normalized to the control (set as unity). Data represent means ± SD from at least three independent experiments.

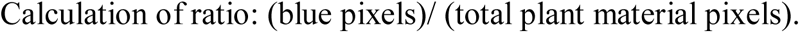

#### 2.8.5 Determination of PAL Enzyme levels

The phenylalanine ammonia-lyase (PAL) activity was assessed following the method outlined by Cheng and Breen (1991). PAL activity was measured by conversion of L-phenylalanine to trans-cinnamic acid. The harpin treated leaf was detached after different intervals, including 2, 4, 12, 24 and 48 hpi. A total of 0.5 g of the leaf tissue was ground in pre-cooled mortar and pestle using liquid-N2 and resuspended into 5 ml of boric acid extraction buffer consisting of 5 mM β-mercaptoethanol, 0.05 M boric acid, 1 mM EDTA, and 0.05 g of polyvinylpyrrolidone. Supernatant was collected after centrifugation at 18,894 rcf for 20 minutes at 4°C. The reaction mixture contained 500 μl of 50 mM boric acid buffer (pH 8.5), 200 μl of 50 mM L-phenylalanine (substrate), 200 μl of crude enzyme extract, 100 μl of distilled water and incubated at 37°C for 1 h. The reaction was stopped by adding 100 μl of TCA (10%). The absorbance (290 nm) was recorded using spectrophotometer. The trans-cinnamic acid molar extinction coefficient (174000 M⁻¹cm⁻¹) was used to calculate PAL activity. The enzyme activity was conveyed as the μM of trans-cinnamic acid formed per minute per gram of fresh weight of leaf (μM/min/g FW).

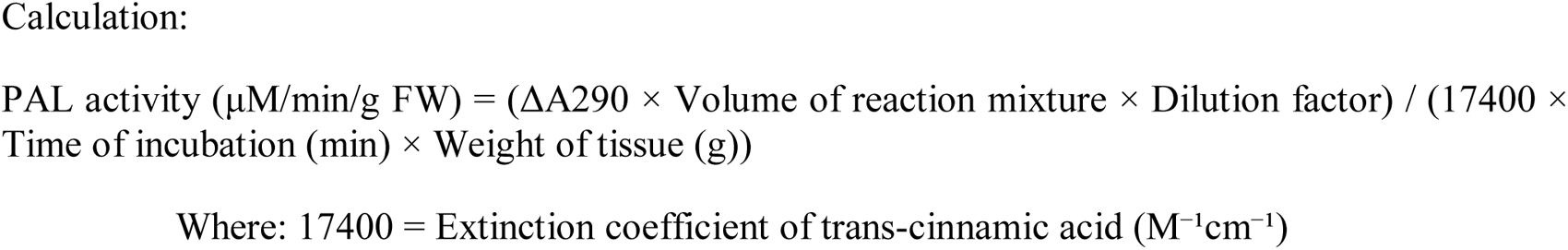

#### 2.8.6 Determination of PPO enzyme levels

The enzymatic activity of PPO was examined by following the method reported by Chen et al., (2022) with few modifications. Catechol, a PPO substrate, was added exogenously based on the protocol. The harpin infiltrated leaf samples were collected at different time intervals (i.e., 3, 6, 12, 24, 48, and 72 hours). The leaf sample (0.5 g) was ground and made into a fine powder using mortar and pestle, and resuspended into a 3 mL extraction buffer (sodium phosphate buffer (100 mM) and 0.1 ml Triton X-100 (0.1% v/v), pH 6). The homogenate was transferred to a pre-chilled tube, incubated for 30 min at 4°C, followed by centrifugation at 18,894 rcf for 30 minutes at 4°C. The supernatant was used as the crude enzyme extract. The reaction mixture contained 1.9 ml of reaction buffer containing 10 mM catechol solution and 0.1ml of enzyme extract supernatant. The absorbance at 420 nm was measured at every 15 seconds for 5 minutes (for a blank, reaction buffer containing 10 mM catechol solution). One unit (U) of PPO activity was defined as the amount of enzyme that causes an increase in absorbance of 0.001 per minute under the assay conditions. PPO activity was determined using the formula:

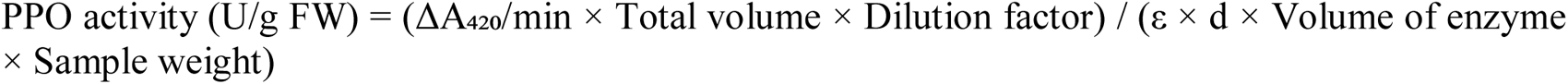

Where: ΔA₄₂₀/min = Change in absorbance per minute, Total volume = Total reaction volume (2 ml), Dilution factor = Any dilution of the enzyme extract, ε = Molar extinction coefficient for quinone formation (24300 M⁻¹cm⁻¹ for benzoquione), d = Path length (1 cm), Volume of enzyme = Volume of extract used (0.1 ml), Sample weight = Fresh weight of leaf tissue (g).

## 3 Results

### 3.1 *In silico* analysis of the primary, secondary and tertiary structures of harpin

#### 3.1.1 Analysis of physicochemical properties of harpin

The amino acid sequence of harpin retrieved from the NCBI was used for the determination of different physicochemical properties of harpin using the ProtParam tool of the ExPASy server (Table 1) (Gasteiger et al., 2005). The theoretical pI and molecular weight of harpin are 4.93 and 34.5 kDa respectively. The aliphatic index was 85.99, and the instability index was 35.97 with a grand average of hydrophobicity (GRAVY) of −0.214. The ProtParam tool also computed the extinction coefficient (Table 1). The sequence contains 19 positively charged residues such as Arg (R) and Lys (K), and 42 negatively charged residues such as Glu (D) and Asp (E).

**TABLE 1.**
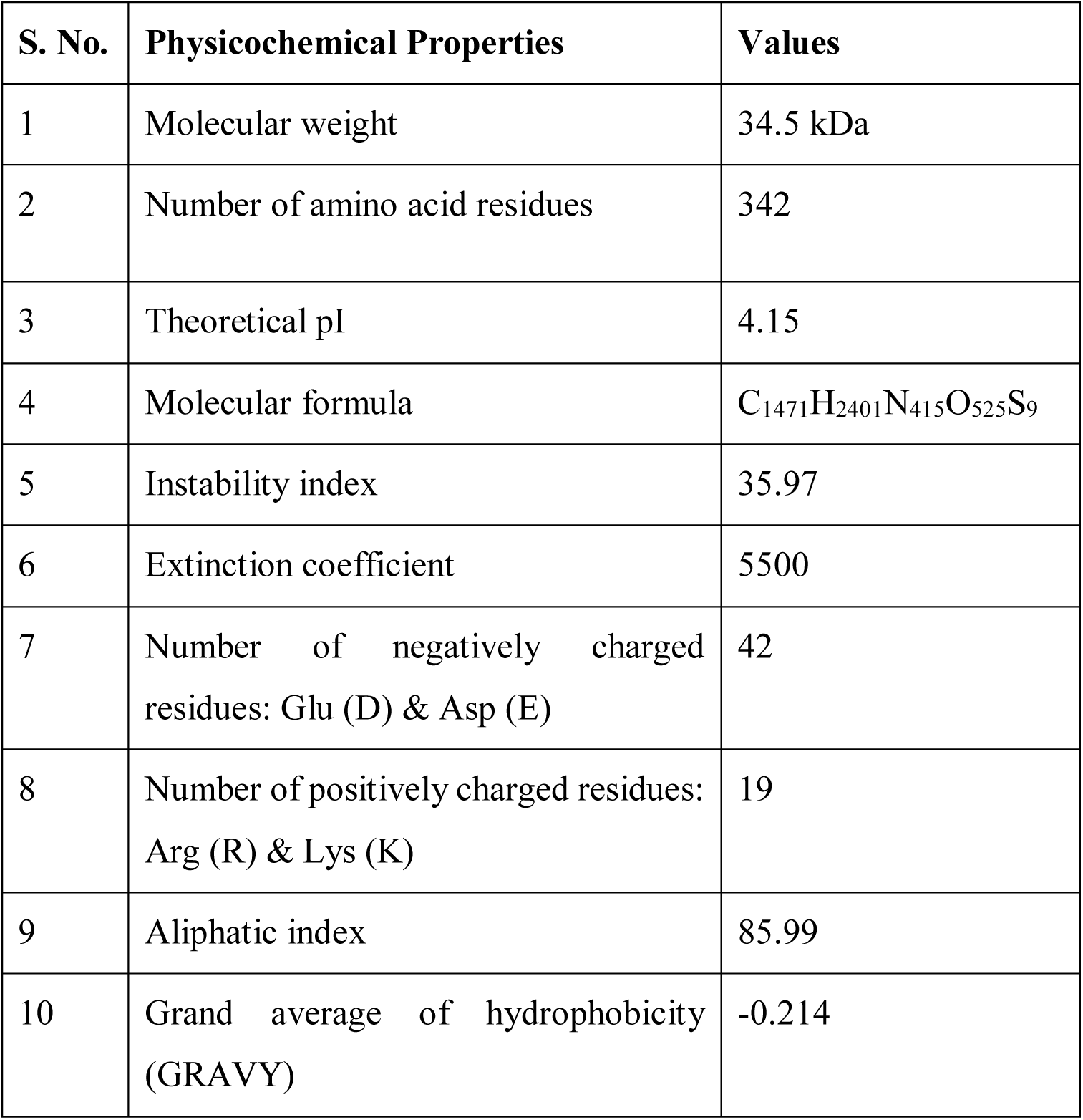
Characterization of harpin (HrpZ2*_Ps_*) using the ProtParam tool.

#### 3.1.2 Secondary structure of harpin

The predicted harpin protein comprised of 50.29% (172 residues) alpha helices, 48.53% (166 residues) random coils and only 1.16% (4 residues) beta sheet. The representative predicted secondary structure of harpin protein is shown in Figure 1A. A total of 42% (144 aa residues) of the total structure was predicted to be of intrinsically disordered regions (Figure 1B) (Jones, 1999).

**FIGURE 1.**
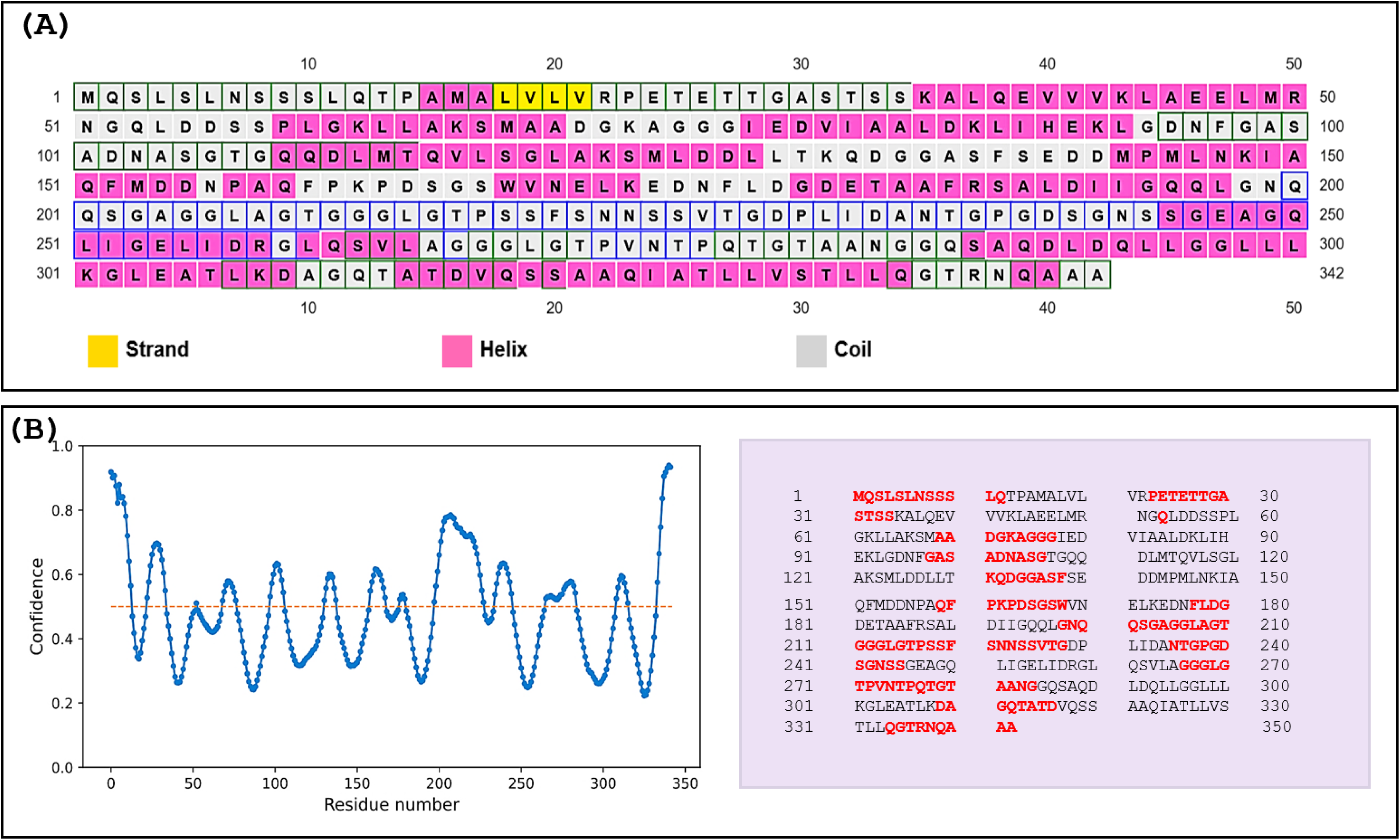
Secondary structure prediction and intrinsic disorder profile of harpin (HrpZ2*_Ps_*). **(A)** The prediction revealed a predominance of α- helical conformations (50.29%, 172 residues shown in pink), followed by random coils (48.53%, 166 residues shown in light colour) and β-sheet architecture (1.16%, 4 residues shown in yellow). **(B)** Left panel: intrinsic disorder propensity plot (blue color) of amino acid residues, with the disordered threshold demarcated by the horizontal yellow dotted line. Right panel: schematic distribution of amino acid residues, where residues exceeding the disorder threshold are highlighted in red color.

#### 3.1.3 Prediction of the tertiary structure of harpin

A 3D model of harpin was generated using the I-TASSER web server and validated with the VERIFY 3D, ERRAT, and PROCHECK tools of the SAVES server, including the Ramachandran Plot analysis (Figure 2). The plot showed 90.1 % of harpin residues reside in the most favored regions, 8.9% are in additionally allowed regions, 0.4 % are in generously allowed regions, while 0.7 % in disallowed areas. The sequence comprises 282 non-proline/non-glycine residues, 46 glycine and 12 proline residues. The 3D model of harpin was visualized using the BIOVIA Discovery studio.

**FIGURE 2.**
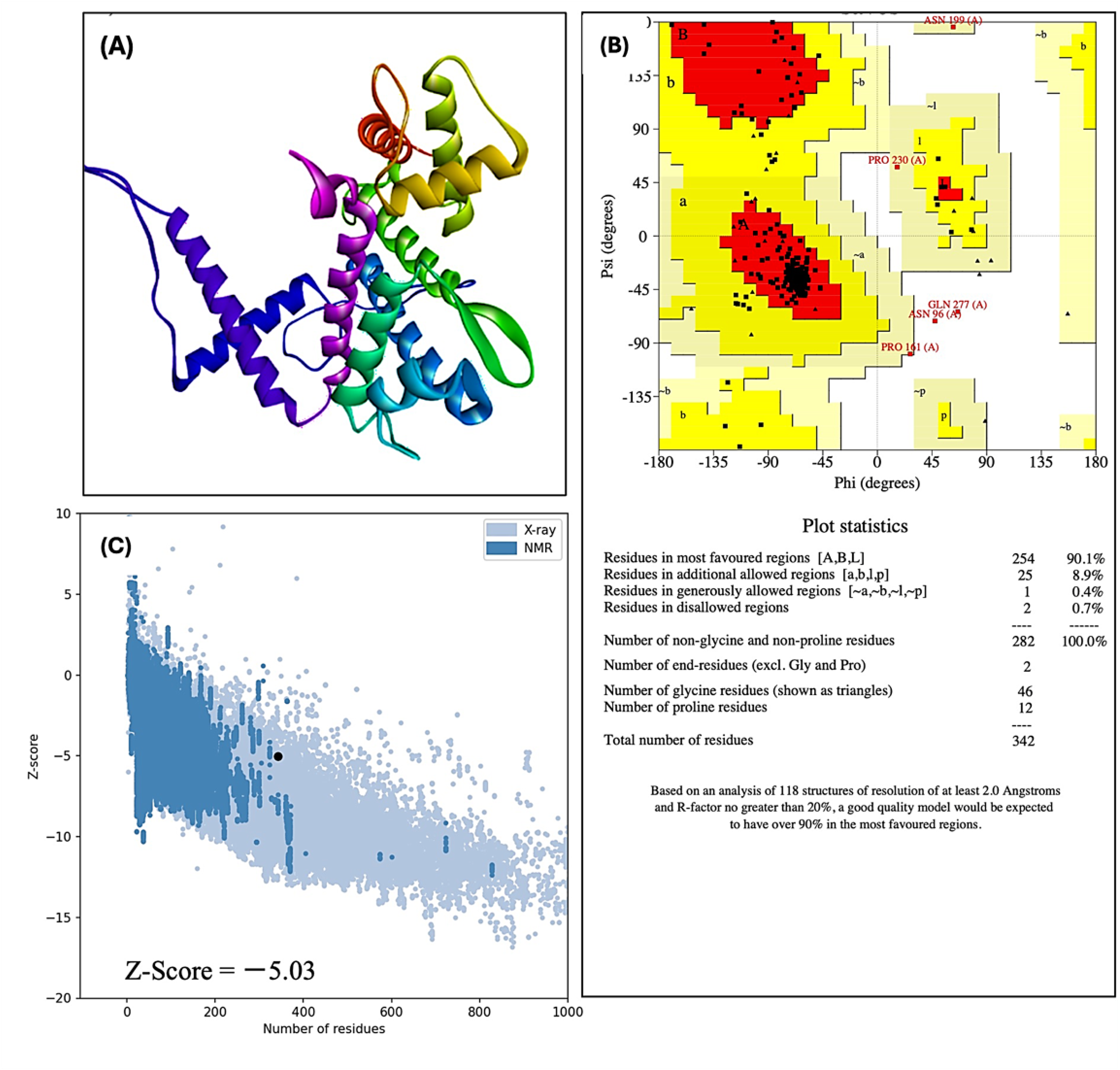
3D modeling and validation of tertiary structure of harpin from *Pseudomonas syringae.* **(A)** 3D model of harpin (HrpZ2*_Ps_*) **(B)** Ramachandran plot with statistical analysis to evaluate the quality of the predicted 3D model. **(C)** Global model quality assessment. The Z-score obtained from the ProSA-web server indicates the overall quality and global structure reliability of the 3D model.

#### 3.1.4 Conserved domain analysis of harpin and transmembrane helix prediction

The mapping of amino acid sequences of the newly identified harpin (HrpZ2*_Ps_*) from *P. syringae* showed that it belongs to the harpin superfamily (Figure 3A). Proteins in this family play various roles, such as eliciting HR in non-host plants and forming a cation permeable pore which plays an essential role in ion-conduction. A signal peptide is present at the N-terminal (from 1 to 23 residues) of the protein as detected by the MEMSAT-SVM method, whereas another region (from 323 to 338 residues) at the C-terminal forms a pore-lining helix, while the residues from 24 to 322, form an extracellular loop confirmed as by the kyte-Doolittle method in TMHMM web server (Figure 3B).

**FIGURE 3.**
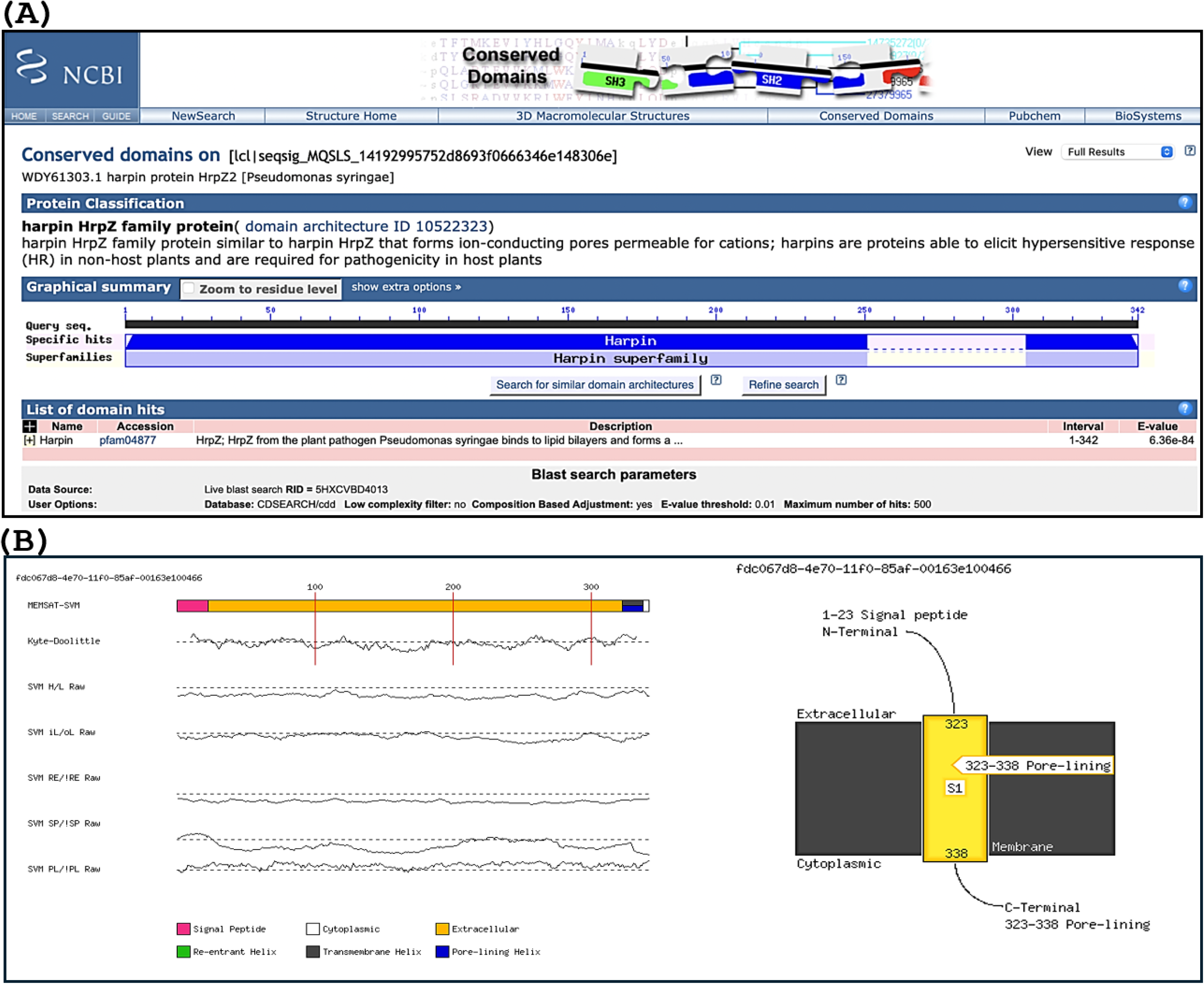
Conserved domain analysis and prediction of transmembrane (TM) helices of harpin (HrpZ2*_Ps_*). **(A)** Conserved domain mapping (indicated by blue shading) reveals specific, high-confidence hits, establishing its homology and putative functional relationship with the harpin superfamily. **(B)** Left panel: transmembrane (TM) helices (color-coded for distinct visualization) of harpin predicted by MEMSAT-SVM tool of the PSIPRED webserver. H/L (Helix/Loop), iL/oL (Inside/Outside Loop), PL (Pore-Lining residue), RE (Re-Entrant helix residue), and SP (Signal Peptide residue). Right Panel: A schematic representation of harpin, with its N-terminus predicted to contain a Signal Peptide (residues 1–23), while a Pore-Lining region (residues 323–338) is identified at the C-terminus.

#### 3.1.5 Comparative domain analysis of harpins

Choi et al., 2013, presented the first domain-based classification of harpins, predicting the secondary structure domain architectures using the PSIPRED web server, and grouping harpins into five classes based on the structural features and similarity. In the present study, we performed a parallel domain analysis of our harpin (HrpZ2*_Ps_*) alongside representatives of each of the five different groups of harpins defined by Choi et al., (2013), incorporating additional input parameters such as the intrinsically disordered regions (IDRs) and residue composition (number of glycines, cystines and leucines) to refine comparisons (Figure 4).

**FIGURE 4.**
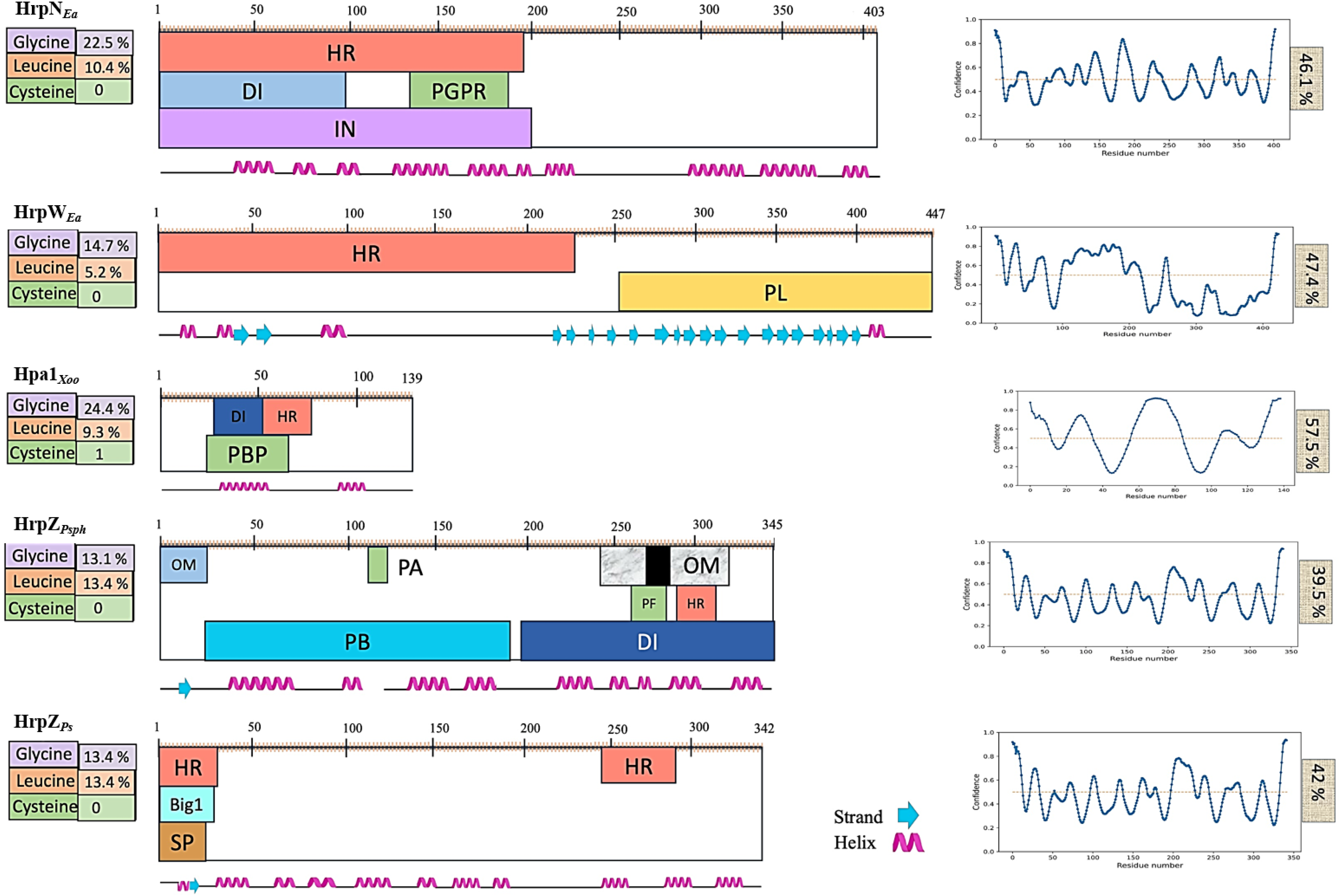
Functional domain architecture of different harpins. Predicted secondary structures (shown in the bottom line), and functional domains are shown for HrpN*_Ea_* and HrpW*_Ea_* from *Erwinia amylovora*, HrpZ1*_Pst_* and HrpZ2*_Ps_* from *Pseudomonas syringae*, and Hpa1*_Xoo_* from *Xanthomonas oryzae*. Information on Glycine-, leucine-, and cysteine-rich regions are displayed on the left, while that of intrinsically disordered regions (IDRs) on the right. HR (hypersensitive response), SP (signal peptide), PL (pectate lyase), IN (interaction with a host protein), OM (oligomerization), PGP (plant growth promotion), DI (defense induction), PA (phosphatidic acid binding), PF (pore formation), and PB (peptide binding).

#### 3.1.6 Subcellular localizations

The subcellular localizations of our harpin (HrpZ2*_Ps_*) predicted using the DeepLocPro-1.0 server is graphically represented in Figure 5 (Moreno et al., 2024). The graph of harpin is leveled with sorting signal importance, which represents the contribution of each amino acid residue of harpin protein to the prediction of its subcellular localization and the peaks of amino acid residues important for corresponding the protein to its predicted location. The subcellular location of the harpin was primarily predicted as extracellular with a high probability (0.9915), showing a strong likelihood that the protein is located outside the cell. At the same time, alternative localizations were also predicted with very low probability. The alternative potential localizations are outer membrane (0.006) and periplasmic space (0.0023), while cell wall and surface, cytoplasmic membrane, and cytoplasmic are predicted to be very low with 0.0001 probability.

**FIGURE 5.**
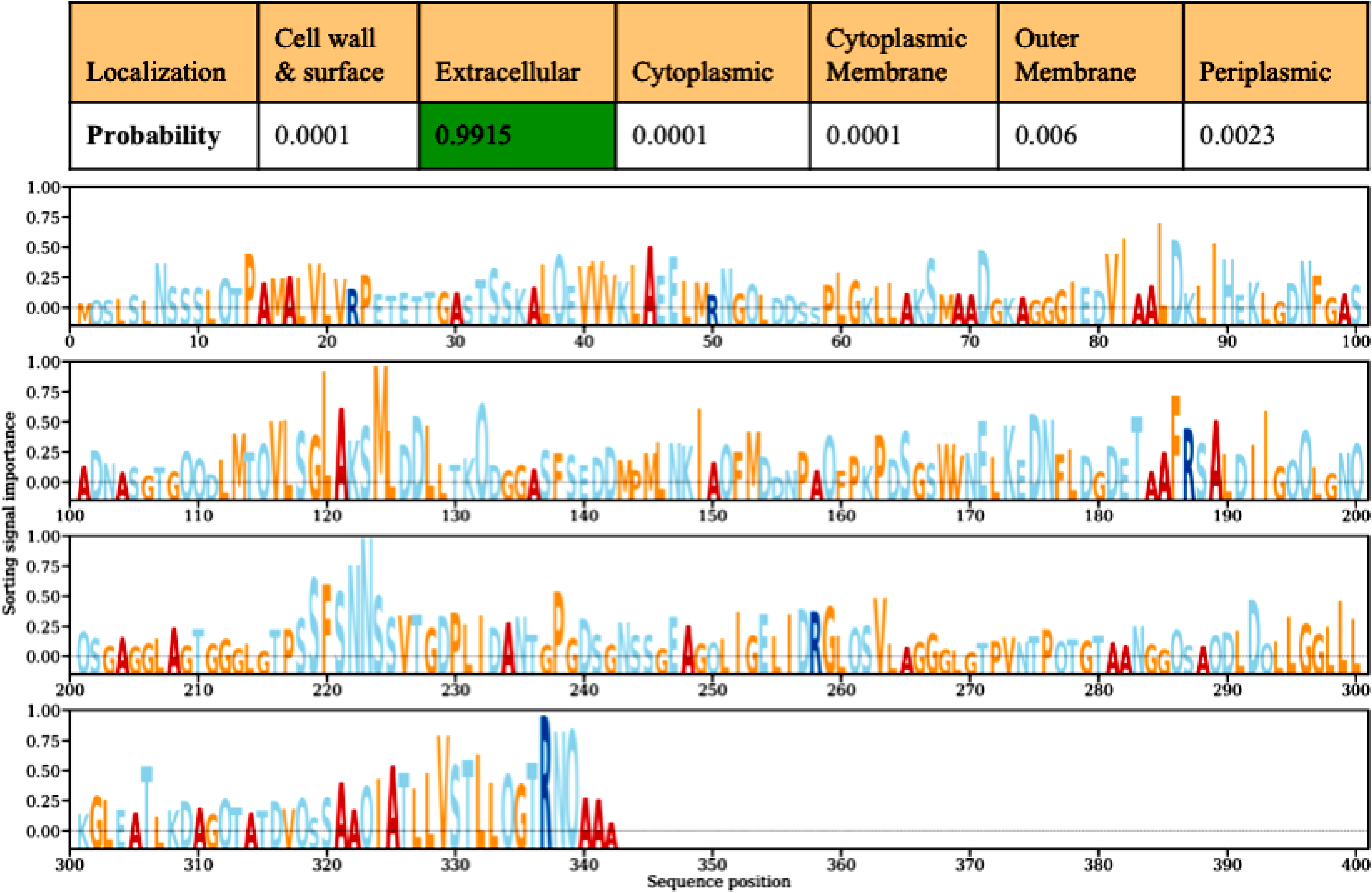
Subcellular localization of the harpin (HrpZ2*_Ps_*) predicted using DeepLocPro-1.0. The top panel shows the highest localization probability of harpin for the extracellular region (0.9915), highlighted in green. The lower panel illustrates the contribution of sorting signals across amino acid residues, representing their relevance to subcellular localization.

#### 3.1.7 Evolutionary study of harpins

Phylogenetic analysis of different harpins showed sequence clustering, which indicates a strong genus-specific grouping, with harpins from the same or closely related species forming tight clades. For instance, the HrpN protein from *E. amylovora* formed an independent clade (highlighted in red), reflecting a conserved evolutionary lineage distinct from other harpins. Similarly, harpins from *P. syringae* pathovars showed broader distributions across several sub-branches (highlighted in green). The new harpin (HrpZ2*_Ps_*) is marked with a red circle in the picture (Figure 6).

**FIGURE 6.**
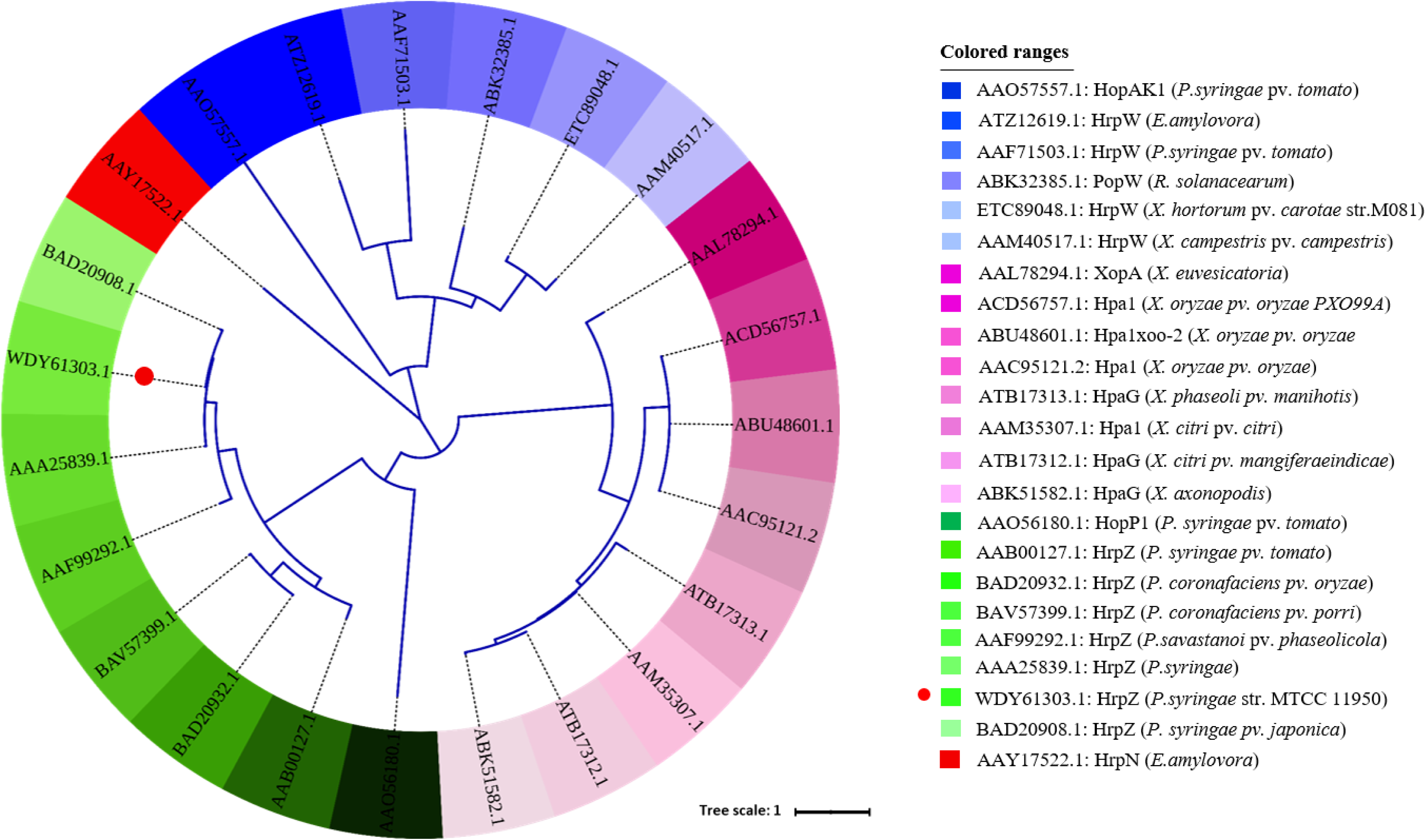
Phylogenetic analysis of 23 different harpins from different bacterial pathogens based on their amino acid sequences. The tree was constructed using MEGA11.0 and visualized with the iTOL tool (https://itol.embl.de/shared/2PuqbCYlUfiSH). Each harpin is color-coded, with respective protein names displayed on the right side.

### 3.2 Purification and western blotting of harpin

The harpin (HrpZ2*_Ps_*) protein was successfully purified from *E. coli* BL21 cells harbouring the harpin gene construct, and the purity was assessed by SDS PAGE (Figure 7), and protein concentration determined by Bradford Assay was 132 μg/ml. Western blot analysis showed specific recognition of His-tagged harpin by an anti-His antibody (Figure 7), confirming successful cloning, expression, and purification of the recombinant protein.

**FIGURE 7.**
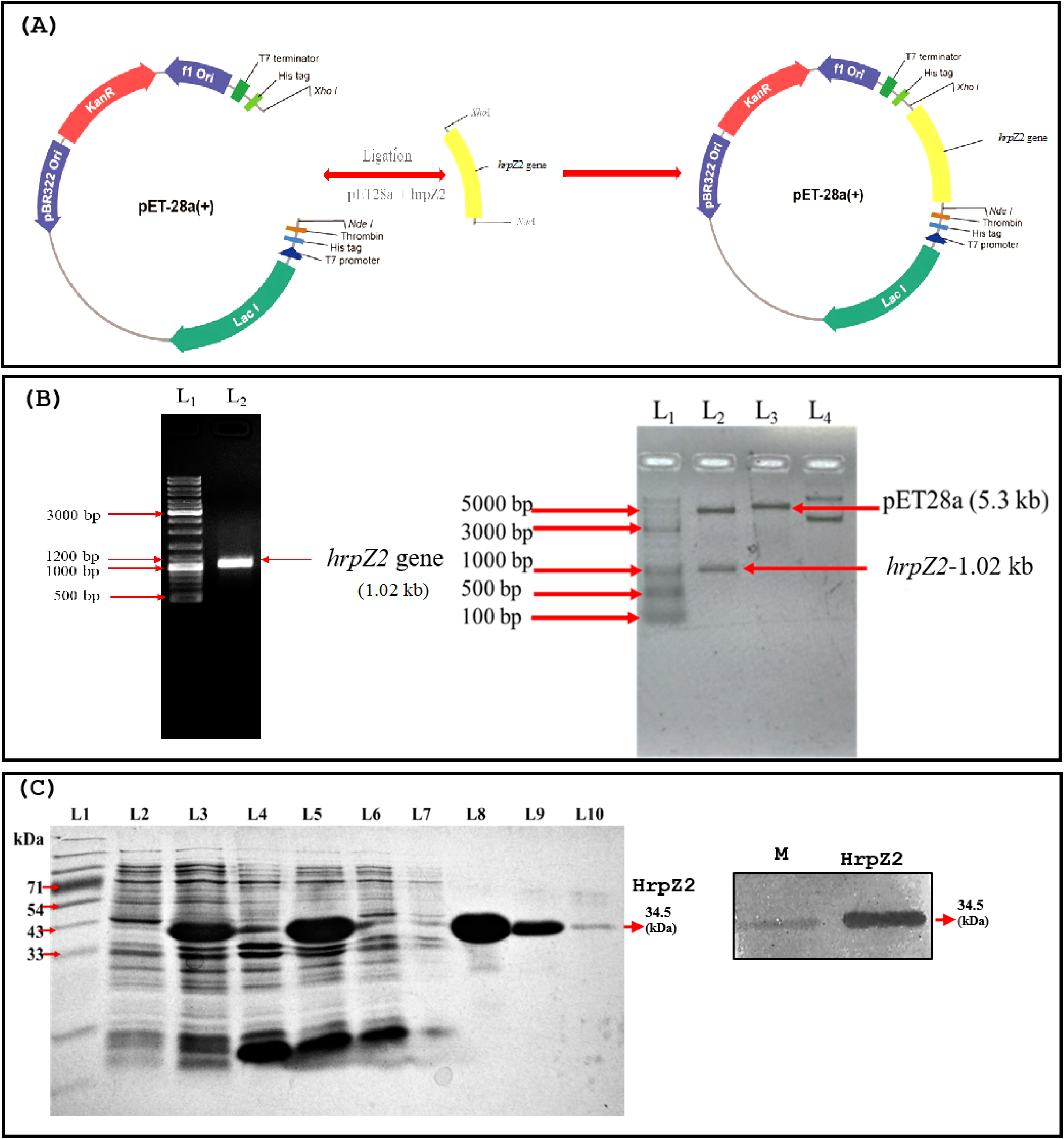
Cloning, SDS-PAGE, and western blot of harpin (HrpZ2*_Ps_*). **(A)** Map of *pET28a-hrpZ2* construct showing cloning of *hrpZ2* at *Nde*I and *Xho*I sites. **(B)** PCR amplification of harpin (Left), and confirmation of cloning of harpin in the construct by double digestion with *Nde*I and *Xho*I (right). **(C)** SDS-PAGE and western blot of harpin. Left panel: SDS-PAGE analysis of harpin expression. L1: protein molecular weight marker; L2 –7: fractions collected during expression and purification steps; L8–10: elution fractions containing purified harpin. Right panel: western blot of harpin immunoblotted with anti-His-tagged primary and secondary (IgG-ALP) antibodies.

### 3.3 Biochemical assays

#### 3.3.1 Determination of range of non-host plants which responds with HR elicitation with harpin

Purified harpin protein was infiltrated into the abaxial (lower) surface of leaves of several non-host plants through a sterile needleless syringe. A clear, visible HR was observed in the localized area of dead cells where the infiltration was done, while buffer treatment showed no such effect (Figure 8A, B). It is reported that elicitor proteins of a critical concentration and above can only induce visible macroscopic HR lesions, and below that concentration, it may fail to induce such visible HR or may induce microscopic lesions (micro-HR) (Peng et al, 2003). This critical concentration may vary from harpin to harpin, plant species variants, methods of application etc. The results from the infiltration experiments with serial dilutions of HrpZ2*_Ps_* revealed that the minimum concentration of the harpin elicitor required to elicit a visible HR was 30 µg/ml (Figure 8E).

**FIGURE 8.**
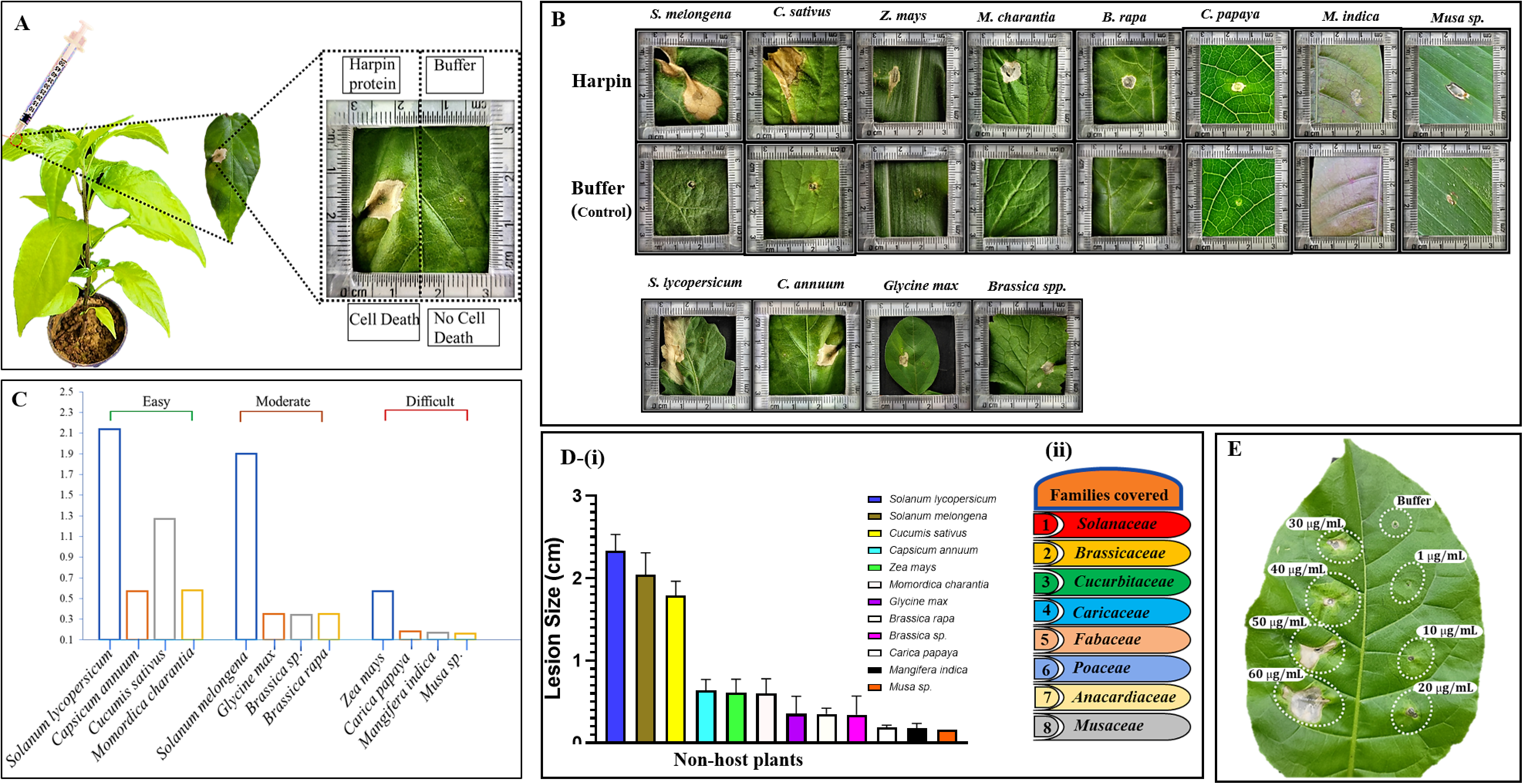
Development of macroscopic hypersensitive response (macro-HR) in non-host plant leaves post harpin infiltration. **(A–B)** HR developed post harpin (50 µg/mL) infiltration on the abaxial surface of leaves, while using sodium phosphate buffer as a control. Photographs were taken 3 days post-infiltration (dpi). **(C)** Categorization of twelve plant species exhibiting different HR responses based on ease of infiltration: easy, moderate, and difficult. **(D)** (i) Bar graph showing diameter (cm) of HR-lesion developed post harpin infiltration on leaves of plants across eight angiosperm families, listed in (ii) Solanaceae (*Solanum lycopersicum, Capsicum annuum, Solanum melongena*), Brassicaceae (*Brassica rapa, Brassica sp*.), Cucurbitaceae (*Cucumis sativus, Momordica charantia*), Caricaceae (*Carica papaya*), Fabaceae (*Glycine max*), Poaceae (*Zea mays*), Anacardiaceae (*Mangifera indica*), and Musaceae (*Musa sp*.). **(E)** Different dilutions of harpin infiltrated into tabocco leaf. Each bar is color coded to correspond to a different plant species. Experiments were independently repeated at least three times, with five leaves used per plant. Data is presented as means ± standard deviation (SD).

To know whether the harpin can elicit HR in various non-host plants, we tested 15 different non-host plants belonging to eight different angiosperm families, including Solanaceae (*Solanum lycopersicum*, *Capsicum annuum* and *Solanum melongena*), Brassicaceae (*Brassica rapa* and *Brassica* sp.), Cucurbitaceae (*Cucumis sativus* and *Momordica charantia*), Caricaceae (*Carica papaya*), Fabaceae (*Glycine max*), Poaceae (*Zea mays*), Anacardiaceae (*Mangifera indica*) and Musaceae (*Musa* sp.). Sodium phosphate buffer (10 mM), pH 7.5 was used as a control. We observed that the onset and intensity of HR varied among the tested plants from different angiosperm families (Figure 8B). Among them, *Momordica charantia* (Cucurbitaceae) exhibited the fastest HR response comparative to other plants with harpin infiltration. In contrast, plants such as *Musa* spp (Musaceae). *Carica papaya* (Caricaceae), and *Mangifera indica* (Anacardiaceae) exhibited a delayed development of macroscopic HR. The bar graph (Figure 8D) represents the relative HR (cell death) development areas induced by infiltration with harpin protein. The above non-host plants are categorized as easy, moderate or difficult on the basis of the ease of infiltration of harpin (Figure 8C).

#### 3.3.2 ROS detection *via* DAB staining and DCFDA fluorescence

To assess ROS accumulation in the harpin-treated leaves, H_2_O_2_ levels were recorded at several time points after infiltration of harpin into tobacco leaves. Harpin triggered rapid H_2_O_2_ accumulation in harpin infiltrated leaves (Figure 9). The level of H_2_O_2_ started increasing by 6 h after infiltration and continued up to 24 h. On the other hand, the level of H_2_O_2_ remained at lower levels consistently in the control leaves. The macroscopic HR was also visible at the site of infiltration after 24 h. The *in vivo* H_2_O_2_ accumulation was also spotted as a brown precipitate using the DAB staining method in the harpin-treated tobacco leaves (Figure 9A) whereas no such brown precipitate was found in the control samples. All these results indicate accumulation of H_2_O_2_ in the harpin-treated tobacco leaves. ROS generated by harpin was also evident and visualized in 6-, 12- and 24-hour post-infiltration (hpi), while observed to be most prominent in a 24-hpi leaf using a ROS-sensitive fluorescent dye 2’,7’-dichlorodihydrofluorescein diacetate (DCFDA). This fluorescent dye detects a broad range of ROS responses, such as generation of H_2_O_2_ and superoxide anion O2-. Tobacco leaves infiltrated with purified harpin showed a strong H_2_O_2_ associated ROS burst, as indicated by increased DCFDA fluorescence, relative to buffer infiltrated controls. In parallel, DAB staining showed a more intense reddish-brown precipitate in harpin treated tissues, further confirming elevated H_2_O_2_ accumulation.

**FIGURE 9.**
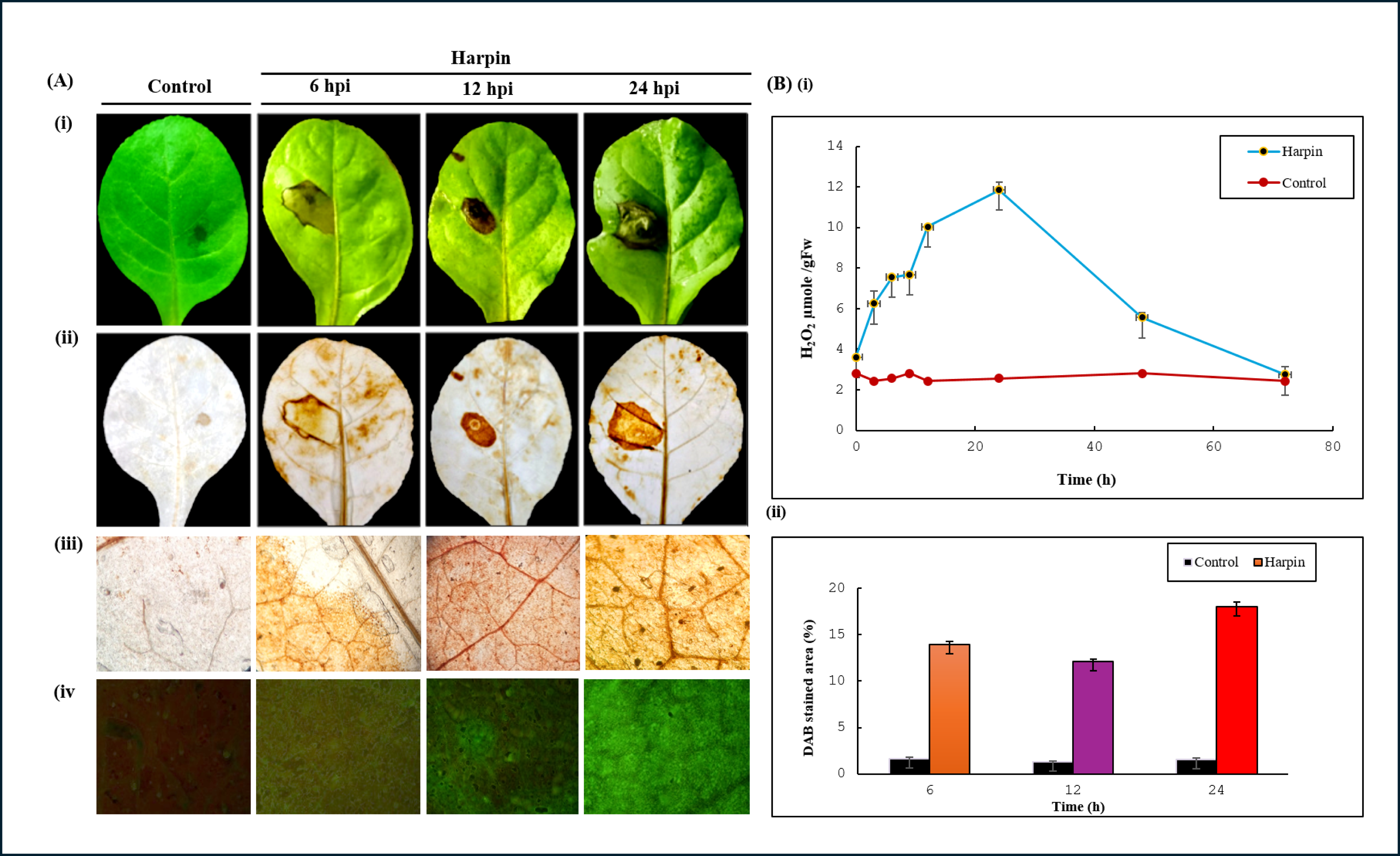
Histochemical detection and quantification of hydrogen peroxide (H₂O₂) in *Nicotiana tabacum* leaves following harpin treatment. **(A)** DAB staining: (i) before bleaching, (ii) after bleaching, (iii) microscopic view, and (iv) DCFDA staining, performed 24 hours post infiltration with harpin (30 µg/mL) or sodium phosphate buffer as control. **(B)** (i) Quantification of H₂O₂ using the FOX reagent method. The x-axis represents H₂O₂ content (µM/gFW). (ii) Measurement of DAB-stained leaf area (%). The x-axis represents DAB-stained area (%). The y-axis denotes time (h) in both graphs.

#### 3.3.3 Quantification of Hydrogen peroxide using FOX reagent method

After infiltration of the purified harpin, a rapid burst of ROS was observed by determining the levels of H_2_O_2_ at various time points. ROS was quantified on tobacco leaves using the FOX reagent method and found that the first peak was visible at 6 hpi (7.55 μM/FW) and then fluctuated till 12 hpi (10.03 μM/FW). The highest peak was observed at 24 hpi (11.85 μM/FW). After that, the level of H_2_O_2_ started declining at 48 hpi (5.55 μM/FW) and at 72 hpi (2.75 μM/FW) in harpin infiltrated leaves. However, the level of H_2_O_2_ was consistently lower in the leaves of control plants. All these results indicate that harpin (HrpZ*_Ps_*) induced the accumulation of H_2_O_2_ in *Nicotiana tabacum* leaves (Figure 9B).

#### 3.3.4 Histological and cytological studies of harpin induced plant defense response

Localized plant cell death is a well-recognized defense response that restricts nutrient availability to the invading pathogens and limits their spread. To detect the harpin (HrpZ*_Ps_*)-induced plant cell death at the site of infiltration, tobacco leaves were infiltrated with the harpin and subsequently stained with trypan blue (Figure. 10A). Harpin treatment clearly induced localized cell death at and around the infiltration site. This defense response is indicative of a hypersensitive reaction of the plant to restrict the pathogen invasion and prevents systemic infection.

**FIGURE 10.**
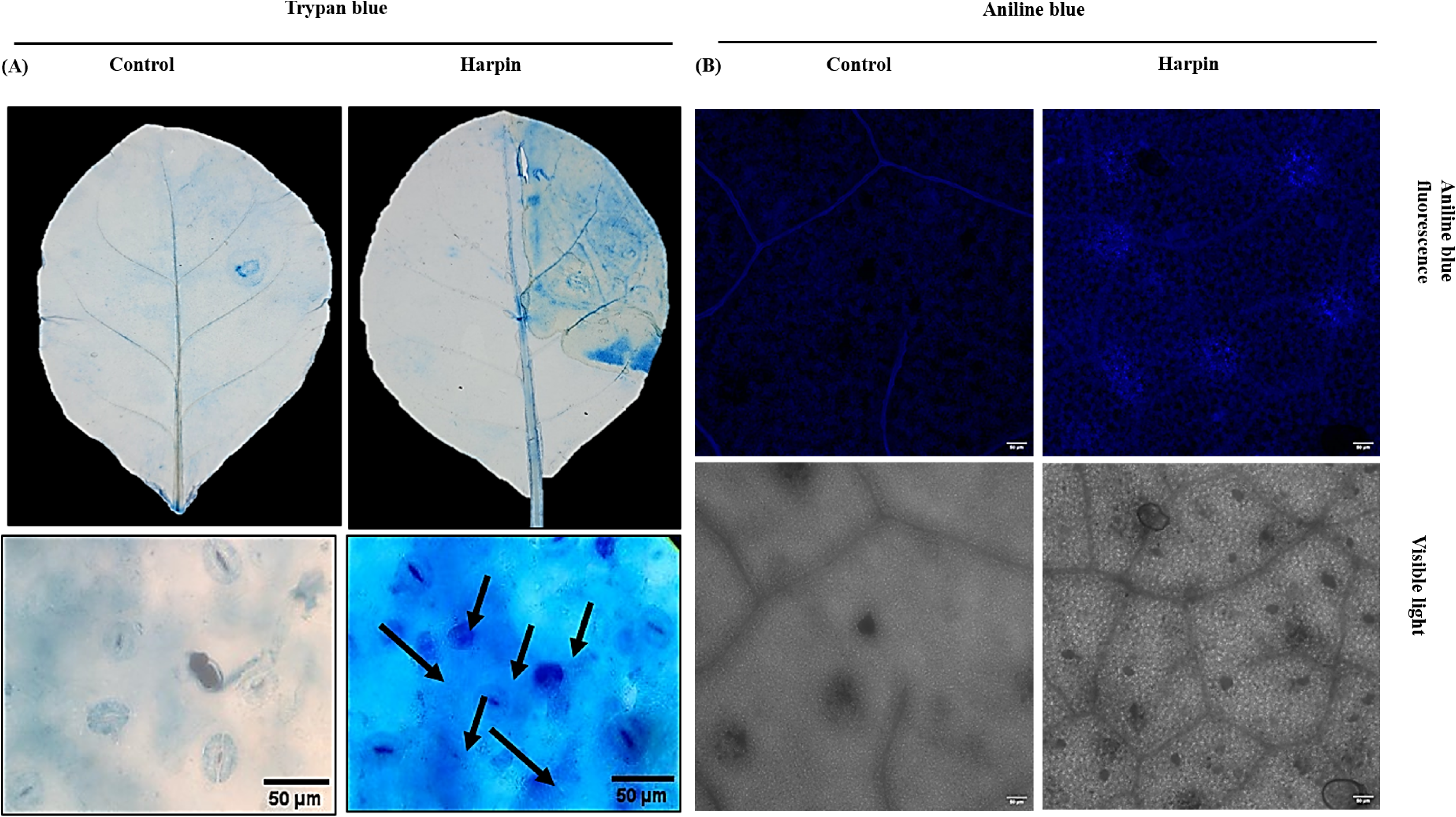
Detection of cell death (trypan blue staining) and callose deposition (aniline blue staining) in *Nicotiana tabacum* leaves post harpin infiltration, through light and fluorescence microscopy. **(A)** Upper panel: leaf infiltrated with sodium phosphate buffer (control-left) or harpin (right). Lower panel: light microscopy images of excised leaf sections from infiltration sites showing hypersensitive response (HR) at 3 dpi. Blue-stained regions (right) (indicated by black arrows) represent HR-induced cell death in harpin-treated leaves, compared to control (left). **(B)** Fluorescence microscopy images showing callose deposition in leaves at 24 hpi. Dark blue patches (right) indicate callose deposition in response to harpin infiltration (30 µg/mL) compared buffer controls (left). Experiments were independently repeated at least three times, with minimum three leaves used per experiment.

Callose is an essential polysaccharide that plays a crucial role in the various processes of plant development and in defense signaling. Callose accumulation was estimated by aniline blue staining of leaf samples followed by observation under confocal microscopy. The images of aniline blue staining of leaves after infiltration are shown in Figure 10B. Quantitative analysis of the callose deposition was done using the number of callose deposits per mm^2^ leaf tissue as shown in Figure 10b. The results show that there were distinctive arrangements of callose accumulation subject on the timepoint that were being observed. While there was no callose deposition observed in the control leaves, the results suggested that harpin induced callose deposition in the infiltrated leaves.

#### 3.3.5 Analysis of plant defense related enzymes in response to harpin

Harpin also elicits defense-related enzymes, notably PAL and PPO, in infiltrated leaves of tobacco. PPO activity was considerably increased in harpin treated leaves compared with control. Results show that the PPO activity began to rise by 6 h after infiltration and continued increasing through 24 h. The activity of PPO in harpin infiltrated leaves was higher than control leaves, with maximum activity observed at 24 hpi (329.21 U/g), after which the level of activity started to decline at 48 and 72 hpi (Figure 11A). Overall, PPO activity remained higher in harpin treated samples than in control leaves.

**FIGURE 11.**
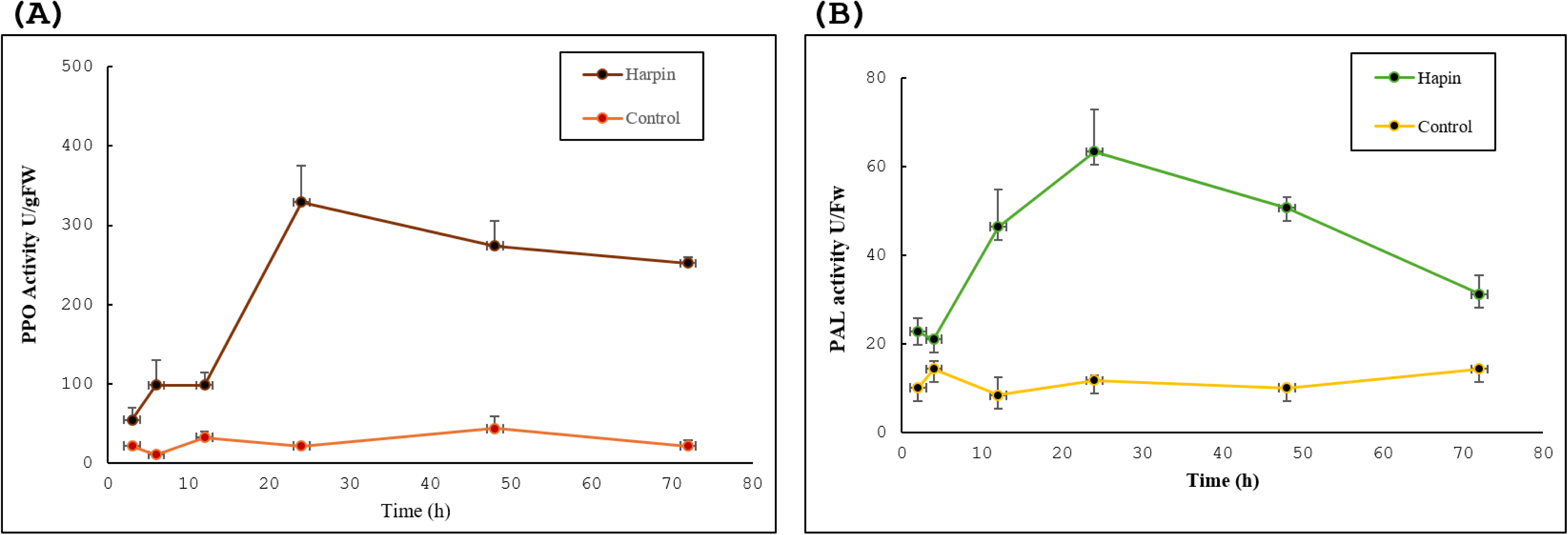
Effect of harpin on induction of defense-related enzyme activities in *Nicotiana tabacum* leaves**. (A)** Polyphenol oxidase (PPO), and **(B)** Phenylalanine ammonia-lyase (PAL) activities increased post harpin (30 µg/mL) infiltration in tobacco leaves, compared to sodium phosphate buffer as control. Both enzymes showed maximum activity around 24 hours post-infiltration. The y-axis represents enzyme activity and x-axis represents time (h) post-infiltration (both graphs). Data is represented as mean ± standard deviations (SD) from three independent experiments, each conducted with three plants.

Our results show another key enzyme, i.e., phenylalanine ammonia lyase (PAL), was also regulated in response to harpin protein in tobacco leaves. PAL activity was moderate initially (21.12 U/g) through 4 hpi, peaked at 24 hpi (63.36 U/g), and declined by 48 hpi (Figure 11B). Overall, PAL activity in harpin-treated leaves was higher than the control.

## 4 Discussion

Plant responses to pathogen attacks are highly complex and entail a multitude of biochemical and cellular processes. One of the key components of plant defense is HR, a form of localized programmed cell death that serves to limit pathogen spread by restricting it to the initial site of infection (Balint-Kurti, 2019). We identified a new harpin elicitor from *P. syringae*, which elicited a hypersensitive response in a wide spectrum of non-host plants. The pairwise protein sequence alignment analysis revealed that this harpin protein (HrpZ2*_Ps_*) is different from the previous harpin (HrpZ*_Pss_*) reported from *P. syringae* pv. *syringae* (He et al., 1993). The total number of amino acid residues in HrpZ2*_Ps_* is 342, while that in HrpZ*_Pss_* is 341 (Figure 1). The major differences observed between these two proteins (HrpZ*_Pss_ vs* HrpZ2*_Ps_*) are listed in Table 2, showing multiple DNA sequence-level mutations (insertions, substitutions, shifts in positions).

**TABLE 2.**
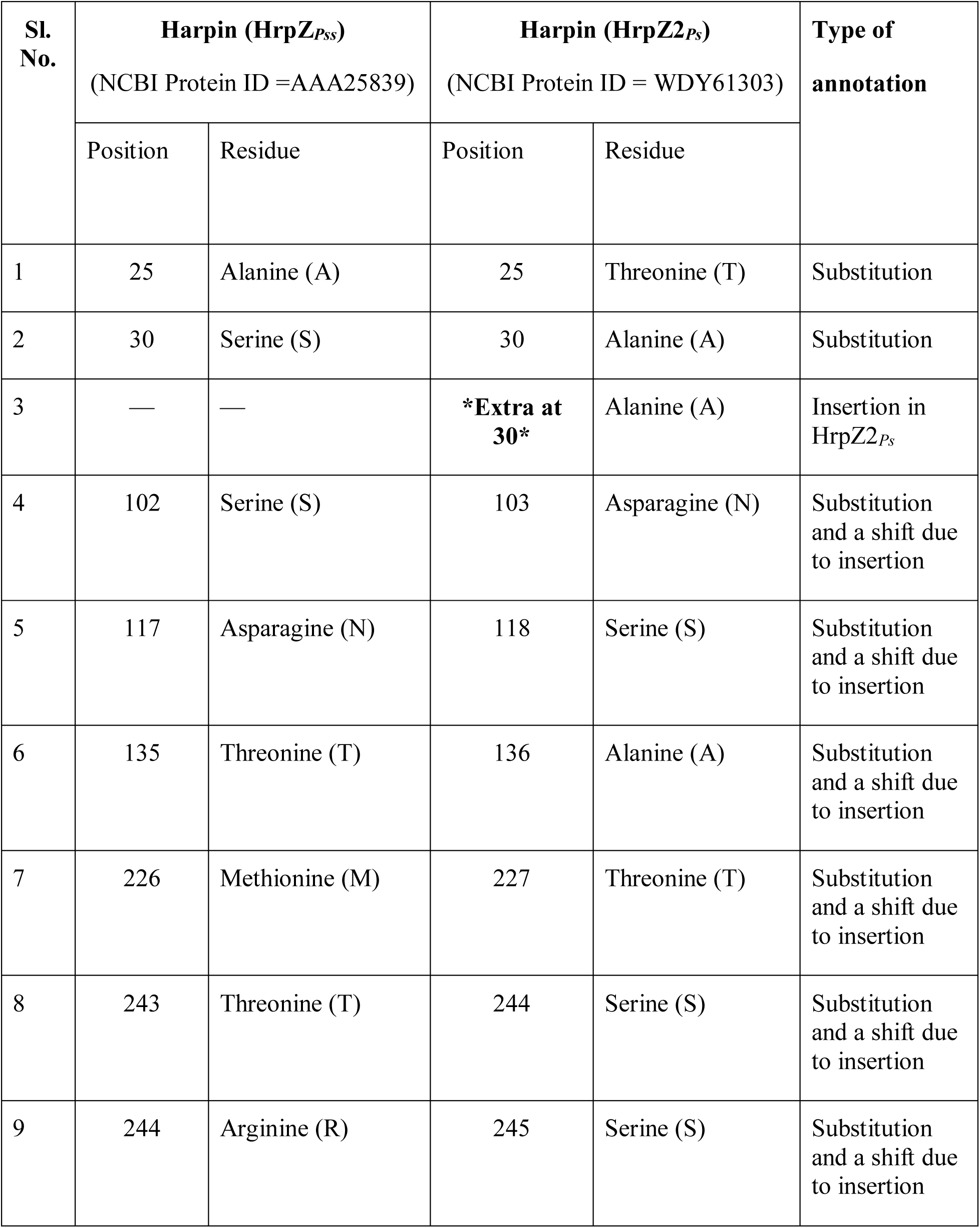
Comparison between HrpZ*_Pss_ vs* HrpZ2*_Ps_* by pairwise protein sequence alignment.

| Sl. No. | Harpin (HrpZ<sub>Pss</sub>)<br>(NCBI Protein ID =AAA25839) |  | Harpin (HrpZ2<sub>Ps</sub>)<br>(NCBI Protein ID = WDY61303) |  | Type of annotation |
| --- | --- | --- | --- | --- | --- |
|  | Position | Residue | Position | Residue |  |
| 1 | 25 | Alanine (A) | 25 | Threonine (T) | Substitution |
| 2 | 30 | Serine (S) | 30 | Alanine (A) | Substitution |
| 3 | — | — | **\*Extra at 30\*** | Alanine (A) | Insertion in HrpZ2<sub>Ps</sub> |
| 4 | 102 | Serine (S) | 103 | Asparagine (N) | Substitution and a shift due to insertion |
| 5 | 117 | Asparagine (N) | 118 | Serine (S) | Substitution and a shift due to insertion |
| 6 | 135 | Threonine (T) | 136 | Alanine (A) | Substitution and a shift due to insertion |
| 7 | 226 | Methionine (M) | 227 | Threonine (T) | Substitution and a shift due to insertion |
| 8 | 243 | Threonine (T) | 244 | Serine (S) | Substitution and a shift due to insertion |
| 9 | 244 | Arginine (R) | 245 | Serine (S) | Substitution and a shift due to insertion |

*In silico* analysis of the physicochemical properties of harpin protein provided valuable insights into its biological properties. HrpZ2*_Ps_* is structurally stable with an instability index of 35.97 (Table 1) and thermally robust. Other characteristics such as hydropathy (GRAVY) and aliphatic index are in accordance with its solubility as well as thermal properties. These characteristics make harpin suitable for the development of elicitor-based plant protection strategies. Based on the analysis of amino acid residues, the theoretical isoelectric point (pI) of the harpin was predicted to be 4.93 (Table 1), suggesting that the protein is acidic.

Analysis of the predicted secondary structure of harpin (HrpZ*_Ps_*) revealed a distinctive structural organization characterized by the presence of alpha helices (50.29 %), followed closely by random coils (48.53 %), with a minimal presence of beta sheets (1.16 %) (Figure 1). The high proportion of coiled regions, comprising nearly half of the total protein structure, indicates a dynamic and flexible architecture that may be essential for conformational adaptability during host-pathogen interactions. Further analysis of the intrinsically disordered regions (IDRs) indicated that nearly 42 % (144 residues) of harpin protein lacks a stable tertiary structure (Figure 1). IDRs are recognized for their role in mediating protein-protein and protein-nucleic acid interactions, primarily where binding adaptability, rapid conformational deviations, and dynamic regulation are needed. The presence of IDRs and their functional importance is well documented in both prokaryotic and eukaryotic systems. Tardigrade, a microscopic eukaryotic animal, has intrinsically disordered proteins (IDPs) and is capable of surviving in extreme environmental circumstances (Boothby et al., 2017). These IDPs contribute to stress tolerance against desiccation and radiation in tardigrades (Yamaguchi et al., 2012). Tardigrade IDPs work through structural plasticity, allowing transient, multivalent interactions critical for their survival under extreme stress conditions. The presence of large IDRs in harpin may offer flexible interactions, adaptability, and the ability to interact with its receptor, components of the plant defense machinery, and promptly trigger plant immune resposes.

Since the initial identification of harpin from *E. amylovora* by Wei et al., (1992), it has been extensively studied for its role in eliciting HR and induction of plant immune responses. Despite decades of work, no crystal structure of harpin has been experimentally determined till date. This lack of structural information hampers mechanistic understanding of its interaction with receptors and downstream immune responses. To address this knowledge gap, a 3D model of harpin protein (HrpZ2*_Ps_*) was predicted by our lab using computational modelling approaches (Figure 2A). Structural validation by Ramachandran plot and ProSA server indicated a well-refined model, revealing 99 % of residues fell in allowed as well as additionally allowed regions (Figure 2B). In contrast, only 0.7 % of residues are in disallowed regions. The Z-score of our computed 3D model of harpin protein as assessed by the ProSA server is −5.03 (Figure 2C), suggesting high stereochemical quality. The overall correctness and reliability of the predicted model support and indicate that the predicted model is reliable. These results support the reliability of the model and, in the absence of an experimental structure, provide a useful framework for probing harpin structure-function relationships and interactions with host immune components (Lal et al., 2023).

The phylogenetic analysis of 23 harpin proteins from different bacterial genera was done. It shows that harpins such as HrpZ1 from *P. syringae* pv. *tabaci*, HrpZ2*_Ps_* from *P. syringae,* and harpins from other *Pseudomonas* species clustered together in a single clade, but with significant intra-genus diversity, suggesting possible functional divergence or adaptation to different host plants (Figure 6). The HrpN protein derived from *E. amylovora* got clustered in another distinct clade (indicated in red) separate from the harpins from *P. syringae* pathovars.

Plant defenses against phytopathogens encompass rapid ROS production and activation of HR (Balint-Kurti, 2019). We investigated H_2_O_2_ accumulation as a defense response triggered by HrpZ2*_Ps_*. In plant defense signaling pathways, H_2_O_2_ functions as a secondary messenger that can drive programmed cell death (Mittler, 2002). Exogenous application or overproduction of H_2_O_2_ can stimulate production of defense-related enzymes and enhance resistance to infection in plants (Mohammadi et al., 2021; Wang et al., 2020). Consistent with this, our DAB staining showed that harpin-induced H_2_O_2_ accumulated locally at infiltration sites, producing a reddish-brown colour signal rather than a uniform leaf-wide pattern (Figure 9). Time course assays showed a parallel increase in DCFDA fluorescence, which increased markedly by 24 hpi relative to buffer-controls, and peaked by 24 hpi, reflecting a strong H_2_O_2_ burst (Figure 9B). Quantification of harpin in tobacco leaves by the FOX assay further confirmed elevated H_2_O_2_ levels upto 9.86 μmol/FW in harpin infiltrated leaves as compared with buffer controls (Figure 9). In line with these ROS dynamics, trypan blue staining demonstrated HR-associated cell death at and around the infiltrated site. Trypan blue staining at 3 dpi showed a deep bluish HR lesion (Figure 10A), reflecting localized H_2_O_2_ buildup and harpin-induced cell death. During the initial stages of bacterial infection, callose deposition reinforces the cell wall as a physical barrier to block pathogen entry. H_2_O_2_ is known to promote callose accumulation (Wu et al., 2018). Harpin infiltration led to higher callose deposition than in controls, with numerous bright light-blue puncta observed by confocal microscopy at 24 hpi, when accumulation was maximal, whereas buffer-treated leaves showed little to no callose accumulation (Figure 10B).

To determine the activities of plant defense-related enzymes (PAL and PPO) at 0- and 24-hpi, tobacco leaves infiltrated with harpin (30 μg/ml) were collected, the enzyme’s activities were measured, and compared with that of buffer as a negative control. Following harpin application, the PPO activity in tobacco increased significantly at 24 hpi, reaching the level of 23.3 U/FW. The activities of PPO in harpin-treated tobacco leaves were found to be higher than those in buffer infiltrated leaves during the entire experimental period (Figure 11A). A rise in PPO activity in leaves treated with harpin suggested that the plant had shown a increased level of disease resistance. Following harpin infiltration, the PAL activity got increased by 24 hpi reaching 32 U/FW (Figure 11B). Consistent with the elevated activities of the two protective enzymes: PAL and PPO, it indicates that harpin boosts key defense enzymes to strengthen plant disease resistance. Similar results of harpin-induced induction of PAL and/or PPO activities were observed by previous reports. Transgenic overexpression of *hrpZm* from *P. syringae* in soybean resulted in increased levels of PAL and PPO, and conferred *Phytophthora* root and stem rot resistance (Du et al., 2018). Harpin application induced the activity of both the PAL and PPO enzymes in the young jujube leaves (Tian et al., 2023). Increased accumulation of anthocyanin pigment was observed in the calluses following harpin treatment, which is due to increased ROS levels and increased PAL activity in callus cells (Daler et al., 2024).

Since the first harpin (HrpN*_Ea_*) was discovered from *E. amylovora* (Wei et al., 1992), it has seen commercial use as a plant growth promoting formulation (MESSENGER®, Eden Bioscience Corporation) in cotton and other crops (https://www.cotton.org/). Harpin-based products have been evaluated by the U.S. Environmental Protection Agency (EPA) with a favorable safety profile for humans and the environment. Names of other existing commercial harpin products include Harp-N-Tek® from Plant Health Care Inc., USA and RX Green Solutions, Axiom Harpin protein from Jensen Distributing, China. In the future, next-generation harpin formulations may offer broad-spectrum disease resistance against various pathogens and pests, and growth promotion in many crops, with strong prospects for commercialization.

In summary, a new harpin (HrpZ2*_Ps_*) from *Pseudomonas syringae*, induced an HR in *Nicotiana tabacum*, a non-host plant, which is associated with various biochemical events such as H_2_O_2_ and ROS production, callose deposition and expression of enzymes involved in defense responses. However, the complete mechanism of action of harpins is not yet unraveled. The results from the present study will add to the knowledge domain of harpins. Further research is needed to comprehensively elucidate the temporal dynamics of H_2_O_2_ accumulation, defense-related enzymes, callose deposition, HR and downstream signaling events in response to different harpin elicitors to elucidate the mode of action of harpins. These findings highlight HrpZ2*_Ps_*’s potential role as an effector that triggers immune responses and lay the groundwork for dissecting its molecular interactions and mechanisms of action.

## 5 Conclusions

A recombinant HrpZ2*_Ps_* protein from *Pseudomonas syringae* was successfully expressed and purified, and its biological activity was confirmed by successful HR induction in *Nicotiana tabacum*. The present study clearly showed that this new harpin (HrpZ2*_Ps_*) caused a significant increase in the activities of two key defense enzymes, PPO and PAL, in *N. tabacum*. Both enzymes showed a time-dependent increase, with notable peak activities at 24 hpi, coinciding with other defense responses such as hydrogen peroxide (H₂O₂) burst, HR cell death, and callose accumulation. The study also demonstrated that harpins can induce HR in a wide range of non-host plants. These findings highlight harpins’ effectiveness in activating plant immune signaling pathways and underscore its potential role in induced resistance strategies for sustainable crop protection. The observed upregulation of PPO and PAL activities supports the view that harpins function as potent elicitors capable of priming plants for enhanced pathogen resistance. Due to the lack of an experimentally determined crystal structure, our validated *in silico* model serves as an essential framework for future functional and interaction studies involving plant receptor proteins or synthetic analogs.

## 7 Author Contributions

KL: Methodology, Investigation, Formal analysis, Visualization, Writing– original draft. AJ: Investigation, Writing – review & editing. VS: Investigation, Writing –review & editing. MM: Data Analysis, Writing– review & editing. PS: Data Analysis, Writing – review & editing. DD: Conceptualization, Investigation, Formal analysis, Data Curation, Visualization, Validation, Funding acquisition, Project administration, Resources, Supervision, Writing - original draft, Writing–review & editing.

## 9 Data Availability Statement

The original contributions presented in the study are included in the article/supplementary material. Further inquiries can be directed to the corresponding author.

